# CNA detection from spatial transcriptomics using SpaCNA

**DOI:** 10.1101/2025.07.08.663688

**Authors:** Zihui Zhang, Xiaochen Wang, Hong Xuan, Zijie Jin, Ruibin Xi

## Abstract

Spatial transcriptomics (ST) enables genome-wide profiling of gene expression while preserving spatial context, yet accurate detection of copy number alterations (CNAs) in tumor ST data remain challenging. Here we present SpaCNA, a spatial-aware computational framework that integrates multi-modal spatial information to improve CNA detection. SpaCNA constructs adjacency graphs using spatial proximity and H&E stained image similarity, refines the raw gene expressions, and implements a Hidden Markov Random Field model for robust CNA state inference. Through extensive benchmarking on simulated data and multi-cancer cohorts, SpaCNA demonstrates superior accuracy and outperforms existing methods in CNA detection and tumor region identification. In real-world applications to breast cancer and colorectal cancer, SpaCNA reveals spatially distinct subclones with context-dependent interactions within the microenvironment. Additionally, applied to a 3D ST dataset of head and neck squamous cell carcinoma, SpaCNA uncovers clone-specific CNAs associated with therapeutic resistance biomarkers across multiple slices. By facilitating precise spatial mapping of CNAs and tumor architecture, SpaCNA significantly enhances our understanding of intratumoral heterogeneity and spatial evolutionary patterns.

## Introduction

Spatial transcriptomics (ST) has emerged as a revolutionary technology in biological research^1,2^. Unlike traditional bulk or single-cell RNA sequencing, which often lose spatial context, it offers a unique ability to profile gene expression while preserving the spatial architecture of tissues^3,4^. In oncology, this capability is particularly valuable for dissecting tumor heterogeneity, mapping distinct cellular niches, and uncovering the spatial organization of the tumor microenvironment^5–7^. By integrating spatial information with transcriptomic data, ST facilitates the identification of region-specific molecular and genomic signatures, spatially coordinated regulatory networks, and key cellular interactions that drive tumor progression and metastasis^8–10^. These insights not only enhance our understanding of cancer biology but also promote the development of precision therapies using spatial information.

Copy number alterations (CNAs) serve as key genomic events in cancer that contribute to tumor evolution and heterogeneity^11,12^. Detecting CNAs from tumor sequencing data is a powerful approach for identifying malignant cell populations and characterizing intratumoral genetic diversity. For single-cell sequencing, well-established computational tools^13–16^ like inferCNV and CopyKAT have demonstrated high accuracy in inferring CNAs. However, methods specifically designed for spatial transcriptomics (ST) datasets remain limited. Compared to scRNA-seq, CNA detection in ST data faces several challenges: (1) ST data often exhibit higher noise levels and sparsity, which can lead to increased false-positive CNA calls; (2) Many widely used ST platforms, such as the 10X Visium, capture transcriptomic data from multiple cells within a single spot, potentially diluting CNA signals in tumor regions and resulting in a higher probability of false negatives; (3) ST data contain multiple modalities of spatial information, including spot coordinates and hematoxylin and eosin (H&E) stained images, making it challenging to effectively integrate these modalities into models for improved CNA inference. Addressing these challenges is crucial for advancing CNA detection in ST data and unlocking its full potential for tumor characterization.

Several studies have attempted to detect CNAs from ST data and have utilized these findings in downstream analyses. For instance, STARCH^17^ assumes that CNAs exhibit spatial continuity and reports results at the cluster level. However, this approach requires the pre-specification of the number of clusters and may be less sensitive to CNAs that are specific to individual or scattered spots. Similarly, CalicoST^18^ employs a cluster-based model to infer allele-specific CNAs at the spot cluster level, but it requires prior single nucleotide polymorphism (SNP) calling from the transcriptome, which is inherently limited due to the sparsity and technical noise of ST data. Additionally, some cancer studies have directly applied methods designed for scRNA-seq, such as inferCNV, to ST data^19^. However, these approaches completely ignore spatial context, which may lead to less accurate results. Collectively, these limitations highlight the need for more specialized computational frameworks that leverage the unique characteristics of ST data to improve CNA inference.

Here, we propose SpaCNA, a novel CNA detection algorithm from ST data. Our method begins by denoising and normalizing the expression profile to obtain initial estimates of copy number ratios. Next, we construct a spot-neighbor graph using spatial information and fit a Hidden Markov Random Field (HMRF) model to assign copy number states. Based on the results from SpaCNA, it also provides functional modules for estimating tumor purity and defining tumor boundary scores, enabling precise delineation of tumor spatial structures. Through extensive benchmarking on simulated datasets and real datasets, SpaCNA outperformed existing CNA detection methods, demonstrating higher accuracy and robustness. When applied to real tumor data, including ST from hepatocellular carcinoma (HCC), breast cancer (BC), and colorectal cancer (CRC), SpaCNA accurately identified CNAs and tumor regions. We also conducted a detailed analysis of the spatial structure of tumors based on CNA results, revealing the unique microenvironments of tumor boundaries and subclones, as well as their potential associations with key CNAs. Finally, we applied SpaCNA to 3D ST data, uncovering insights into tumor biology and progression along new dimension. These findings highlight the potential of SpaCNA as a powerful tool for enhancing our understanding of tumor heterogeneity and spatial organization.

## Results

### Overview of SpaCNA pipeline

SpaCNA takes as input a gene-by-spot expression matrix with spot coordinate and the H&E stained image. The SpaCNA workflow is organized into three modules: gene expression enhancement, chromosome segmentation, and inference of copy number states (Figure 1A and **Methods**). Compared to single-cell RNA-seq data, the additional spatial information of ST data can provide information of the neighboring relationships and histological features of spots. SpaCNA creates an adjacency graph of spots based on their coordinates and the similarity of H&E stained image tiles. Adjacent spots in the graph are more likely to display similar expressions and thus, the copy number states. Due to the high noise and strong sparsity inherent in spatial transcriptomic data, SpaCNA smooths the gene expression by fitting a first-order dynamic linear model for each spot, and followed by averaging neighboring spots using the adjacency graph. SpaCNA then segments the genome by dividing chromosomes into non-overlapping bins of 1 million base pairs (Mbps) and aggregates gene-level expression to the bin level. Using non-tumor cells identified by SpaCNA or user provided as a baseline, SpaCNA estimates the copy number ratio for each bin and applies the BIC-seq^20^ algorithm to jointly infer potential chromosomal breakpoints across all spots. Finally, to achieve results with enhanced interpretability, we treat the continuous copy number ratios as observed values and regarded the underlying discrete copy number states as latent states, thereby constructing a statistical model. To further incorporate spatial similarity, SpaCNA establishes a HMRF model using the spot adjacency graph and iteratively optimizes the likelihood function to assign copy number states to each chromosomal segment. We demonstrated the effects of each step using a breast cancer sample. The incorporation of histological features aids in filtering out the edges between different regions in the adjacency graph (Supplementary Figure 1A), while gene expression enhancement and the HMRF model reduce the noise in the copy number ratio and the final copy number state, respectively (Supplementary Figure 1B, C).

**Figure 1.**
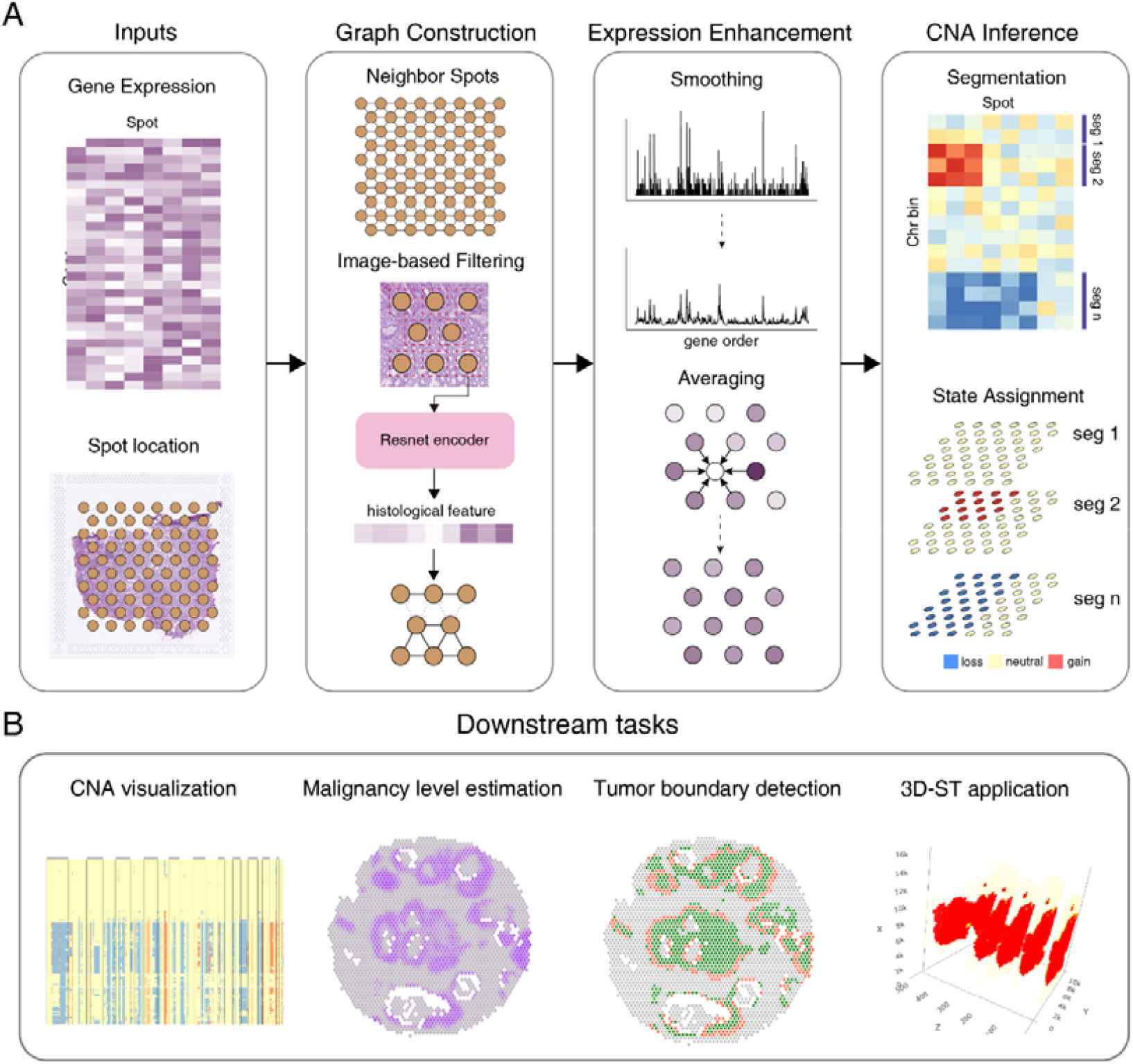
Overview of SpaCNA workflow. **(A)** SpaCNA takes gene expression and spatial information as inputs. First, SpaCNA constructs spot adjacency graph and enhance raw expressions. Then, SpaCNA preforms chromosome segmentations and adopts HRMF model to assign copy number states for each segment. **(B)** SpaCNA enables a suite of downstream tasks including heatmap visualization, malignancy level estimation, boundary score calculation and application to 3D data.

Building on the inferred copy number states, we developed a suite of downstream analytical modules to enhance SpaCNA’s utility (Fig. 1B and **Methods**). We provide the intuitive visualization tool using heatmaps, allowing for easy identification of recurrent CNA events. For tumor identification, we designed a regression model to estimate the malignancy level within each spot. In addition, to better explore the spatial structures, we defined a boundary entropy score to quantify the CNA changes among adjacent spots. As demonstrated in the following analyses, these modules significantly enhance the effectiveness and interpretability of SpaCNA in characterizing tumor genomic landscapes.

### SpaCNA shows superior performance in simulation study

We utilized simulation datasets to evaluate the performance of SpaCNA against other read-depth based CNA methods, including STARCH^17^, CopyKat^16^ and inferCNV^15^ (**Methods**). In brief, we arranged spots in a square grid with three spatially clustered subclones, each harboring a combination of unique CNAs and shared CNAs. To simulate CNAs, we modeled the expression distribution of each gene from a non-tumor reference data, and scaled the distribution means by multiplying the copy number ratio to generate expression values. We applied CNA detection methods on these datasets and assessed their performance using sensitivity and precision. We designed two scenarios to mimic the different type of real datasets: (1) high-purity tumor tissue, where tumor spots contain no non-tumor cells, and (2) mixture tumor tissue, where tumor spots include both tumor and non-tumor cells. To ensure a more comprehensive evaluation, we further generated simulated datasets under various configurations by modulating noise levels, dropout rates, and the number of CNAs.

Overall, SpaCNA showed higher levels of precisions and recalls than other methods (Figure 2). In the high-purity tumor tissue scenario, our method consistently achieved precision and recall values above 0.95 across all configurations (Figure 2A). Although higher dropout rates posed challenges due to increased data sparsity, the overall F1 scores of our method remained robust. InferCNV also demonstrated strong performance, except for slightly lower recall, which could be attributed to its lack of spatial information integration (Supplementary Figure 2). This limitation made it less effective at detecting certain CNAs when the signal was weak. STARCH showed unstable performance, especially in high dropout rate conditions, primarily due to its reliance on spots clustering. As noise levels increased, clustering errors became more frequent, resulting in significant deviations in the final outcome. CopyKAT had the lowest F1 scores among all conditions and scenarios, as it only reported continuous copy number ratios. To make a fair comparison, we manually converted the results into discrete copy number states using a fixed threshold, which reduced the method’s accuracy and flexibility. In the mixture tumor tissue scenario, the performance of all algorithms was reduced, while SpaCNA still outperformed others (Figure 2B). In this scenario, the CNA signals were diluted, causing a significant decrease in recalls across all methods, especially when the proportion of tumor cells is low. The precisions of SpaCNA remained consistent, while all three other methods had a slight to significant drop. This validated the robustness of SpaCNA and highlighted the necessity of the expression enhancement step, which leveraged spatial information to aggregate similar spots and amplify the diluted CNA signals. In addition, we also evaluated the accuracy of subclone clustering from the detected CNA profile (Supplementary Figure 3), where our method again demonstrated the best performance.

**Figure 2.**
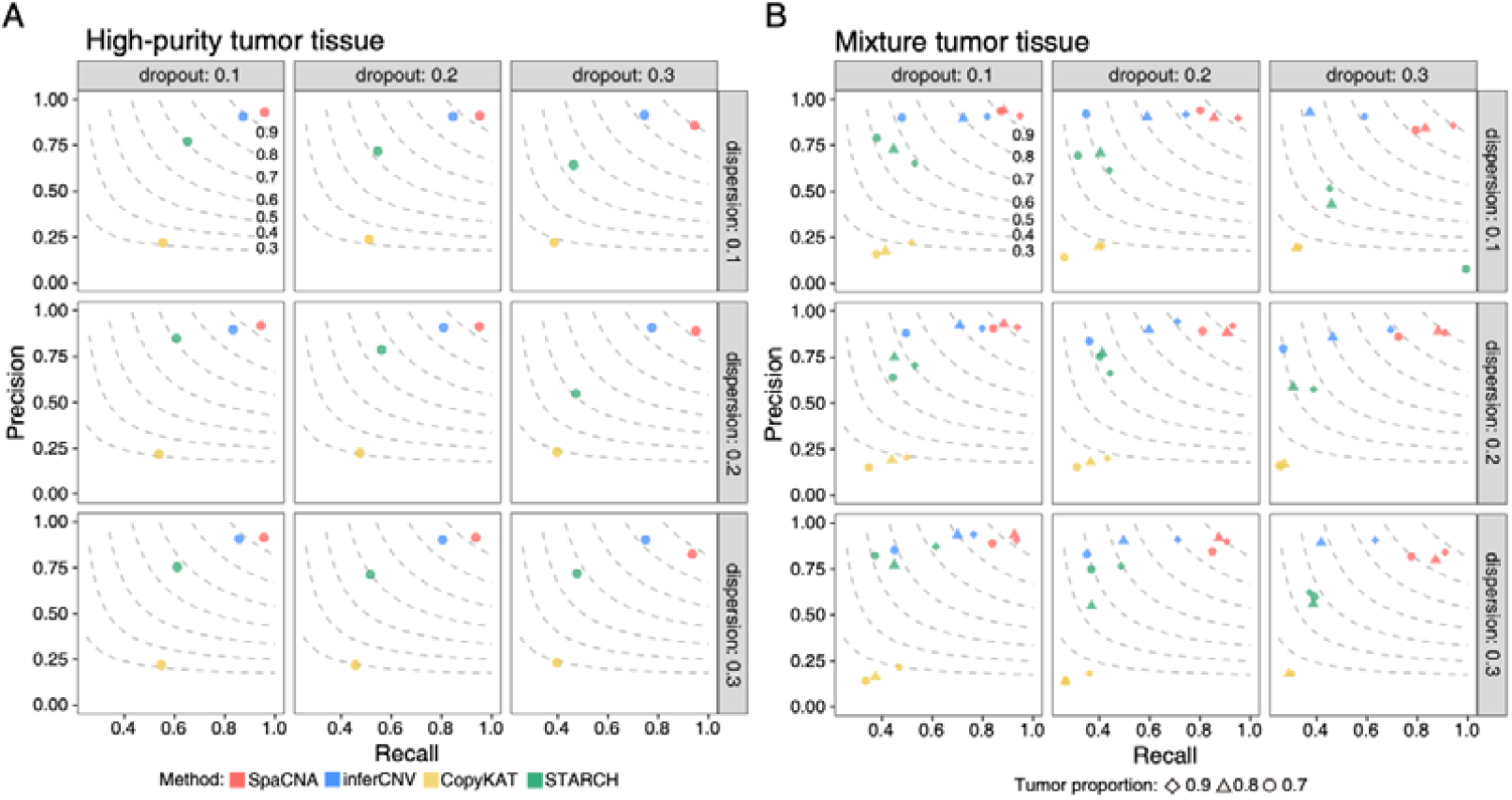
Performance on the simulation study. **(A, B)** The precisions and recalls of SpaCNA and other methods under various dropout and dispersion conditions in the high-purity tumor tissue datasets **(A)** and mixture tumor tissue datasets **(B)**. The shape in figure B represents the different tumor cell proportions in the simulation settings. The dashed lines are the contour lines with constant F1-scores (F1-scores are marked in the top-left figure).

### SpaCNA accurately identifies key CNAs in real tumor data

We further applied SpaCNA and other methods to multiple real spatial transcriptomics datasets. To evaluate the accuracy of each method, we first analyzed a cohort of liver cancer patient samples that included both spatial transcriptomics and whole-exome sequencing (WES) data^21^. This dataset comprised samples from patients with hepatocellular carcinoma (HCC), intrahepatic cholangiocarcinoma (ICC), and combined hepatocellular-cholangiocarcinoma (cHC), covering both the tumor leading edge regions and tumor core regions (Supplementary Tables 1). We used FACETS^22^ to infer bulk-level copy numbers from the WES data as the gold standard. We aggregated spot-level CNAs predictions into bulk level by calculating the modes for each bin, then compared these CNAs with the gold standard to evaluate the performance using precision, recall, and F1 score (**Methods**). SpaCNA outperformed other methods, achieving the overall highest F1 scores (Figure 3A). When precision and recall were evaluated separately, SpaCNA also remained among the top performers (Supplementary Tables 2 and 3). We speculated that some slices like cHC1-L may not perfectly match its WES data, since the consensus CNAs called from ST data (e.g. 1p amplification) were missed in WES (Supplementary Figure 4), suggesting possible limitations when considering WES as gold standard.

**Figure 3.**
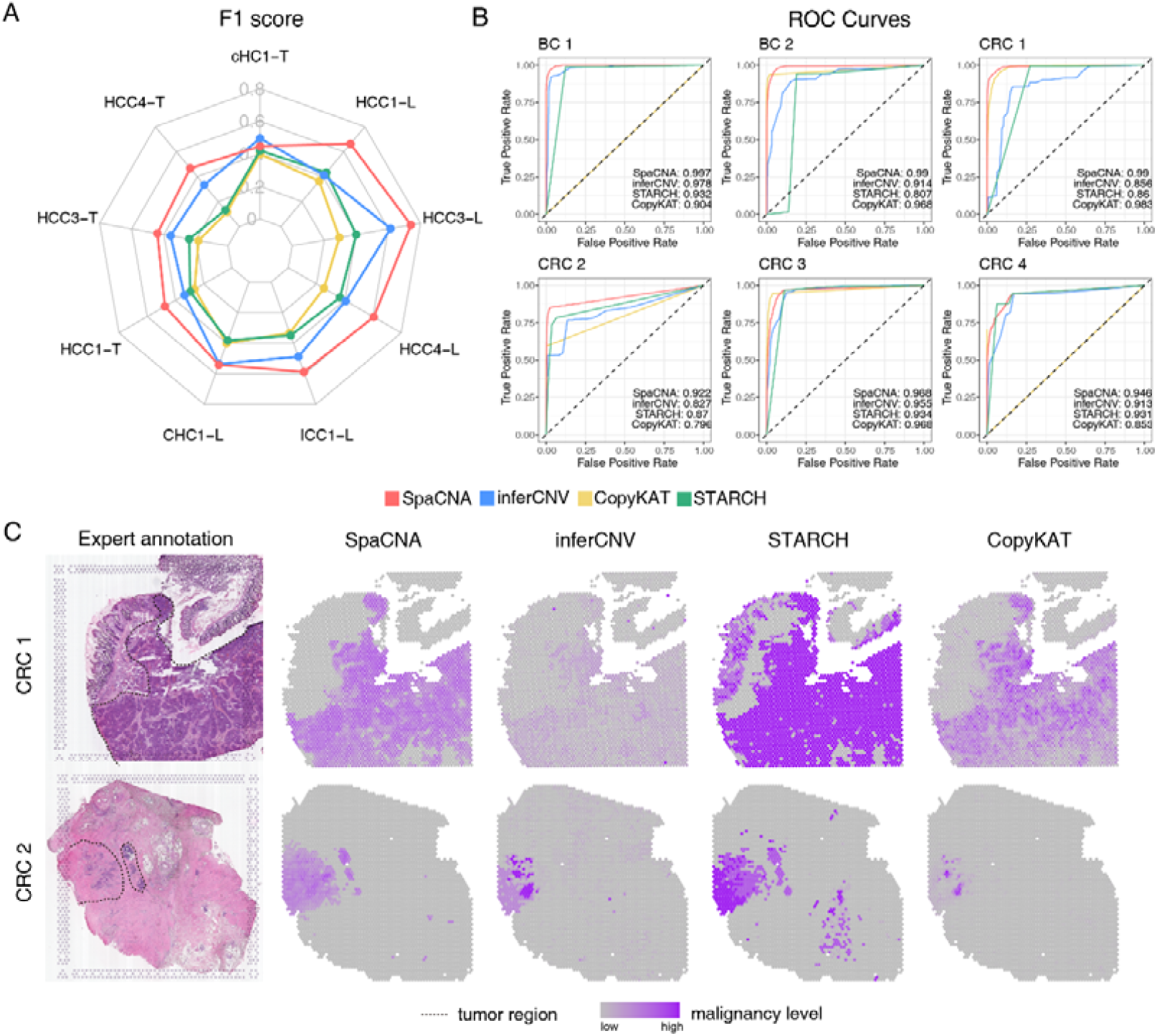
Performance on real-world tumor datasets. **(A)** The F1-scores of SpaCNA and other methods in nine liver cancer datasets. The grey polygons are the contour lines with constant F1-scores (F1-scores are marked on the lines). **(B)** The ROC curves for tumor region identification by SpaCNA and other methods in six datasets. The AUC values of each method are marked in the lower right. **(C)** The expert-annotated tumor regions (first column) and the malignancy scores calculated from CNA profiles in CRC 1 and CRC 2.

Next, we examined the ability that methods accurately identified the spots carrying CNAs. Practically, CNAs were often used as the metric to distinguish tumor from non-tumor regions, where tumor regions usually harbored more CNAs than non-tumor regions. We additionally collected data from breast cancer and colorectal cancer to elucidate the tumor detection ability of SpaCNA and other algorithms. We utilized deconvolution method SpaCET^23^ to infer the tumor proportion per spot and delineated tumor and normal spots accordingly for each sample as the ground truth (**Methods**). Based on the CNA results from each method, we calculated the malignancy score for each spot and evaluated the accuracy of tumor region identification using the binary classification metric ROC AUC (**Methods**). SpaCNA achieved an overall highest AUC score across the samples (Figure 3B and Supplementary Figure 5A), indicating that the CNAs inferred by SpaCNA were mostly located within tumor regions, while minimizing false positives in normal regions. This aligned with the fact that CNAs more frequently occurred within the tumor regions. In contrast, other methods were less stable and exhibited lower AUCs in several samples. For instance, STARCH showed low AUC in BC 2, CRC 1, and HCC-1L; inferCNV performed poorly in CRC 1, CRC 2, and HCC3-L; and CopyKAT underperformed in CRC 2 and CRC 4. To further validate these findings, we calculated Pearson’s correlation between the malignancy scores and tumor proportions, with SpaCNA achieving the highest correlations in most samples (Supplementary Figure 5B). Additionally, for BC 1, BC 2, CRC 1, and CRC 2, tumor regions were annotated based on histological knowledge from H&E stained images by pathology experts (Figure 3C and Supplementary Figure 5C). While all methods correctly detected tumor regions in BC 1 and BC 2, STARCH erroneously identified CNAs in the non-tumor upper-left region of CRC 1, and CopyKAT and inferCNV failed to detect the striped tumor region in CRC 2. Overall, these results strongly validated SpaCNA as a highly effective and reliable method for CNA calling and tumor region identification based on ST data, demonstrating its superiority over existing approaches.

### SpaCNA effectively elucidates the spatial structure of the tumor

In the field of tumor spatial transcriptomics, CNAs and functional modules provided by SpaCNA offer advantages for analyzing tumor spatial structures. For a breast ductal carcinoma sample (referred to as BC 1 in the previous section), we employed boundary scores to distinguish the tumor core region from the tumor boundary. H&E stained images exhibited a strong concordance between the identified boundary spots and histological structures (Figure 4A, B). To investigate the tumor microenvironment, we utilized a breast cancer single-cell reference dataset and applied the deconvolution method cell2location^24^ to estimate cellular composition within each spot. Subsequently, we analyzed the variations in cell type distribution across the boundary region, non-tumor region, and tumor core region (**Methods**). Furthermore, we conducted Gene Set Variation Analysis (GSVA)^25^ to identify differentially enriched cancer hallmark pathways (**Methods**). We found that the tumor boundary was closely linked to enhanced cell cycle activity. Specifically, compared to the tumor core region, pathways associated with E2F and G2M checkpoints were highly expressed in the boundary region (Figure 4C, D), both of which play key roles in regulating cell proliferation. Additionally, we observed an enrichment of cycling tumor cells and cycling perivascular-like (PVL) cells in this boundary region (Figure 4E). These findings suggest that the cells at the tumor boundary are undergoing rapid proliferation and active cell cycle regulation, potentially playing a crucial role in tumor progression and invasion.

**Figure 4.**
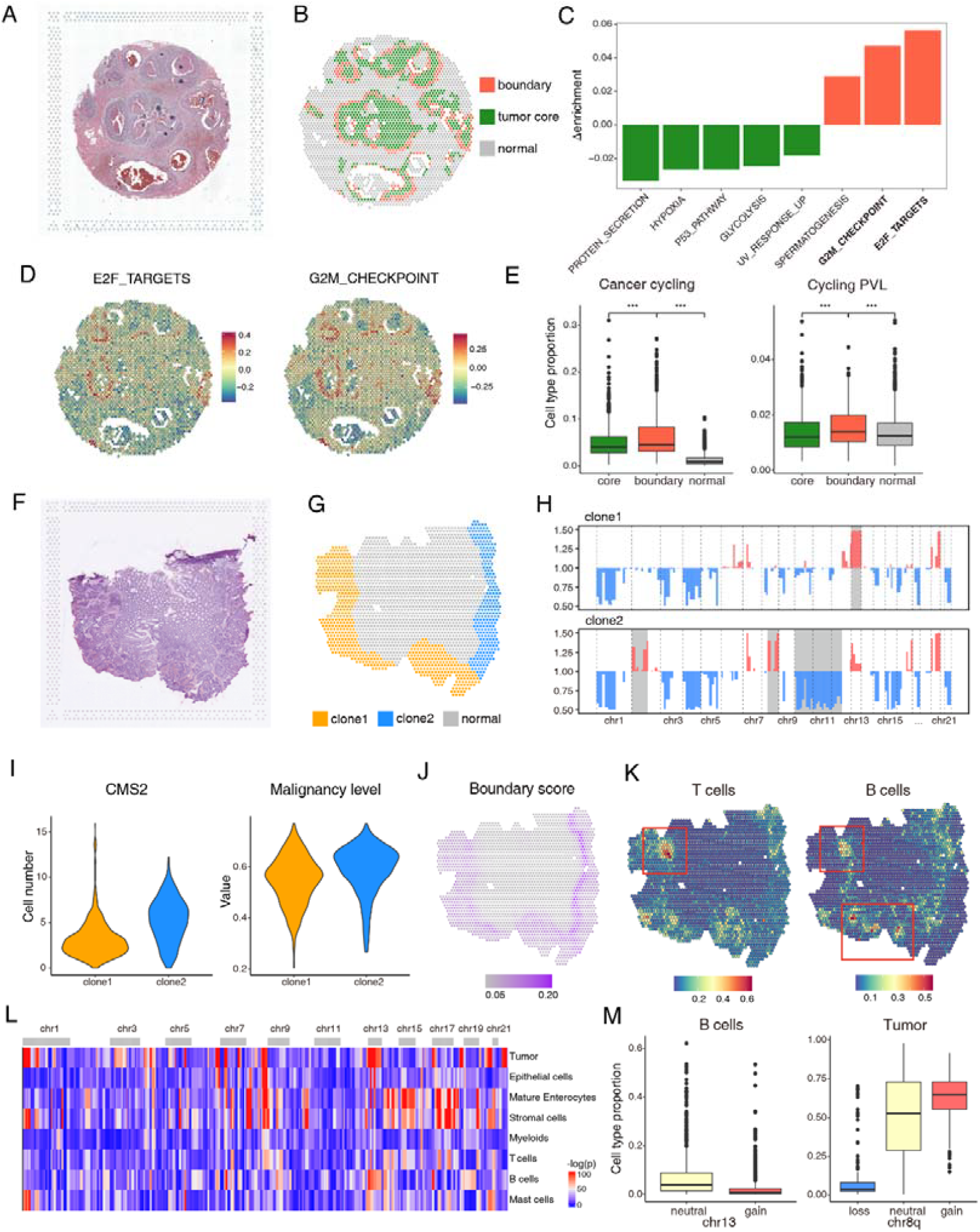
SpaCNA elucidates the spatial structure of tumors. **(A)** The H&E stained image of sample BC 1. **(B)** The tumor core regions, tumor boundary regions, and the non-tumor regions classified by SpaCNA. **(C)** Differential enriched cancer hallmark pathways between tumor boundary regions and tumor core regions. **(D)** The enrichment scores of pathway E2F_TARGETS and G2M_CHECKPOINT. **(E)** Boxplots showing the proportion of cycling tumor cells and cycling perivascular-like (PVL) cells in different regions. The asterisks indicate a significant difference in distribution (Wilcoxon’s rank sum test, P < 0.001). **(F)** The H&E stained image of sample CRC 1. **(G)** The location of two tumor subclones identified by SpaCNA. **(H)** The mean CNA profile of two subclones. The grey background indicates regions with significant CNA difference. **(I)** Violin plots of consensus molecular subtype 2 (CMS2) cell numbers and the malignancy scores in two subclones. **(J)** The boundary scores calculated by SpaCNA. **(K)** The spatial distribution of T cells and B cells, with clusters highlighted by red boxes. **(L)** Analysis results of association between cell type proportions and CNA states. P values are obtained from Kruskal-Wallis tests. **(M)** Boxplots showing the proportion of specific cell types (left: B cells; right: tumor) in spots with different copy number states (left: copy number neutral and gain in chr 13; right: copy number loss, neutral and gain in chr 8q).

In the colorectal cancer sample CRC 1 (Figure 4F), we identified two distinct tumor subclones based on the results from SpaCNA. These subclones, designated as S1 and S2, were situated on the left and right sides of the slice, respectively, exhibiting significant CNA differences on chromosomes 8, 9, and 13 (Figure 4G, H). We further analyzed these two clones in terms of cell type composition and tumor-associated pathways. Notably, spots in S2 displayed higher malignancy level and were predominantly composed of consensus molecular subtype 2 (CMS2) cells (Figure 4I). GSVA results indicated that the several cancer hallmark pathways were more enriched in S2 (Supplementary Figure 6A), suggesting a higher degree of malignancy in this subclone. Additionally, we found significant differences at the boundaries of the two clones. The boundary score surrounding S2 was significantly elevated (Figure 4J), whereas clone S1 exhibited a less distinguishable boundary, indicating tumor cells in S1 had more frequent interactions with the adjacent non-tumor tissue. This observation was supported by the distribution of immune cells, where an accumulation of immune populations, including T cells and B cells, was observed in the boundary regions of S1 (Figure 4K). Moreover, previous research has indicated that the CMS2 cell, which is enriched in S2, typically demonstrates lower levels of immune infiltration. Together, these findings suggest that S1 is an immune-active subclone, while S2 appears to be more immunologically isolated.

The role of CNAs may be critical in shaping the tumor microenvironment. In this slice, clone S2 exhibited amplification on chr 8q, where the key oncogene *MYC* located. Both clones exhibited elevated *MYC* expression compared to the normal regions, with S2 exhibiting significantly higher expression (Supplementary Figure 6B), suggesting that increased copy number on chr 8q may lead to *MYC* upregulation. The MYC_TARGET_V2 pathway also exhibited a similar pattern (Supplementary Figure 6C). To further explore the association between CNAs and cell types, we conducted a comprehensive analysis of multiple colorectal cancer slices. Specifically, we examined whether the proportions of different cell types varied across spots with distinct CNA states. Our findings revealed associations between several chromosomal segments, including chr 1p, chr 8q, and chr 13, and the abundance of tumor cells and B cells (Figure 4L). For instance, in regions exhibiting amplification on chr 13, we observed a decrease in the presence of B cells. Conversely, in spots where chr 8q losses occurred, there were fewer tumor cells, while regions with chr 8q gains showed an increased presence of tumor cells (Figure 4M). These trends were consistent with the distinctions observed between clones S1 and S2, highlighting the potential impact of CNAs on cellular composition within the tumor microenvironment.

### SpaCNA reveals 3D CNA landscape

In recent years, researchers have advanced the field of spatial transcriptomics by extending it from two-dimensional (2D) to three-dimensional (3D) contexts, which has greatly improved our understanding of tumor evolutionary patterns. With minimal adjustments, our method can also be effectively applied to the data produced by these 3D technologies. In this section, we utilized the human head and neck squamous cell carcinoma (HNSCC) dataset generated by Open-ST^26^ and employed SpaCNA to uncover its CNA landscape. This dataset consists of 19 consecutive slices, capturing the transcriptomes of over one million cells in total. To optimize computational efficiency, we merged the cells into approximately 20,000 pseudo-spots (**Methods**). Then we adapted the adjacency graph construction step in SpaCNA to facilitate connections between adjacent slices, resulting in detection outcomes that maintain 3D consistency.

As a result, SpaCNA achieves consistency across different slices and effectively reveals the continuous 3D structure of the tumor (Figure 5A). The tumor can be spatially divided into three subclones (referred as S1, S2, and S3), which exhibit distinct CNA profiles as revealed by SpaCNA. In all three subclones, SpaCNA identified common CNAs including gains of chromosome 7p and losses of chromosome 3p (Figure 5B, C). Notably, several CNA regions encompass oncogenes linked to tumor progression, such as *EGFR* on chromosome 7 and *CCND1* and *CTTN* on chromosome 11. SpaCNA also identified clone-specific CNAs, particularly a copy number gain on chromosome 3q that was unique to S1 (Figure 5B, C). This CNA segment harbors key cancer-related genes such as *PODXL2* and *PIK3CA*, which are known to play an important role in HNSCC^27,28^. However, other CNA detection methods failed to characterize this alteration (Supplementary Figure 7), possibly due to the smaller size of this subclone.

**Figure 5.**
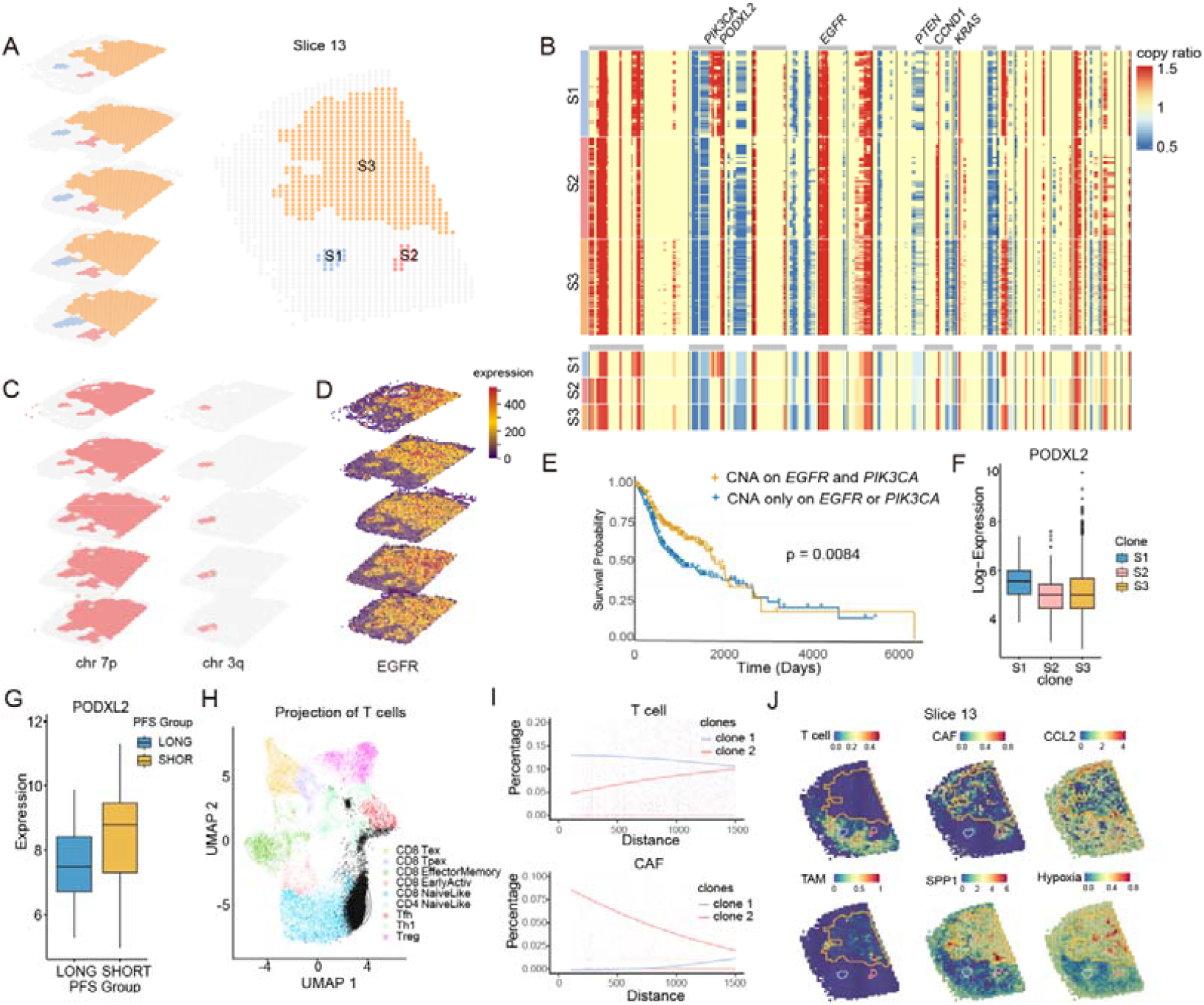
The application of SpaCNA on a 3D ST dataset. **(A)** 3D visualization of tumor clones in slices 14, 15, 16, 17, and 18, and 2D visualization in slice 13. **(B)** The CNA heatmap of three tumor subclones (top; due to the large number of spots in S3, 300 were randomly selected for the plot) and the average CNA profile of each clone (bottom). **(C)** Spots with copy number gain on chr 7p (left) and chr 3q (right). **(D)** Gene expression pattern of *EGFR* in five sections, which co-localized with the chr 7p copy number gain. **(E)** The survival analysis of TCGA samples harboring copy number gains in both *EGFR* and *PIK3CA*, versus those with either *EGFR* gains or *PIK3CA* gains. **(F)** Boxplots of *PODXL2* expressions in three tumor subclones. **(G)** Boxplots of *PODXL2* expressions in patient groups with long PFS and short PFS. **(H)** UMAP plot of a reference T cell dataset and the mapped ST spots (black dots). **(I)** The cell abundance of T cells and CAFs around S1 and S2. The scatter plot indicates the distance and cell proportion for the spots, and the lines were fitted by averaging the cell proportions at specific distance and smoothing in a distance window. **(J)** The spatial distribution of T cell proportion, CAF cell proportion, *CCL2* gene expression, TAM cell proportion, *SPP1* gene expression, and hypoxia scores in slice 13.

The genes located within CNA regions likely have a close connection to tumor progression and prognosis. For example, *EGFR* located within a copy number amplified region and is highly expressed across the overall tumor area (Figure 5D). Additionally, we used COMMOT^29^ to estimate the ligand-receptor interaction strength (**Methods**) and observed that ligands associated with *EGFR*, such as *TGFA*-*EGFR* and *HBEGF*-*EGFR*, exhibited higher intensities in the tumor region (Supplementary Figure 8). Given the critical role of *EGFR* in tumor growth and progression^30^, as well as the proven clinical efficacy of agents like cetuximab^31^ (a monoclonal antibody that specifically binds to *EGFR*), adopting an *EGFR* blockade strategy for this sample may be an effective treatment option. Within the S1-specific amplification on chromosome 3q, there are also genes that are related to tumor prognosis. Numerous studies have reported that amplification of this region serves as a tumor driver and is associated with higher recurrence rates, increased mortality, and shorter overall survival. Genes on 3q, such as *ECT2, PRKCI*, and *PIK3CA*, form an oncogenic cassette that promotes a cancer stem cell-like phenotype and interacts with *EGFR* and *KRAS* to drive tumor progression. *PIK3CA* is also reported to drive tumor growth signaling pathways and is closely associated with tumor progression and poor prognosis. Based on the TCGA cohort (**Methods**), we found that patients with CNAs in both *EGFR* and *PIK3CA* exhibited worse survival than those with CNAs on either *EGFR* or *PIK3CA* region (p=0.0084, Figure 5E). *PODXL2* is another gene located on chromosome 3q, and it serves as a resistance biomarker for the commonly used *EGFR*-targeted therapy cetuximab. Previous studies have found that among patients treated with cetuximab, *PODXL2* expression was higher in those with short Progression-Free Survival (PFS) compared to those with long PFS^32^ (Figure 5F, G). These findings suggested that the S1 remained at risk for further malignant progression.

The three subclones exhibited distinct profiles in both CNAs and tumor microenvironments. T cells were predominantly localized around S1 and S2 without infiltrating into the tumor region (Figure 5I, J). Mapping of the original single-cell level T cells using ProjecTILs^33^ (**Methods**) suggested that the majority of these T cells exhibited a naïve T cell phenotype (Figure 5H), which are typically enriched in lymph node. Notably, the lower-left corner region corresponds to the original lymph node, and S1 was located within the lymph node, suggesting that tumor cells had invaded the lymphatic system. In contrast, S2 and S3 were enriched by cancer-associated fibroblasts (CAFs), with S2 being completely encapsulated by CAFs. We observed that regions with elevated *CCL2* expression partially overlapped with CAF-enriched areas, particularly around S2 and S3 (Figure 5I, J). As *CCL2* promotes the transition from fibroblasts to CAFs via *CCR2* and facilitates the recruitment of myeloid-derived suppression cells (MDSCs), these findings suggested the presence of an immune exclusion microenvironment in tumor subclone S2 and S3. In addition, tumor-associated Macrophages (TAMs) were observed in S3, accompanied by higher expression of *SPP1* (Figure 5J). Previous studies have shown that the presence of TAMs along with elevated *SPP1* levels can induce hypoxia in tumor regions. Accordingly, we observed elevated hypoxia scores and *VEGFA* gene expression in S3 (Figure 5J), indicating a hypoxic and angiogenic microenvironment. These findings highlight the utility of SpaCNA in resolving CNA-defined subclones and their corresponding microenvironmental contexts. The integrated analysis of genomic and spatial transcriptomic features offers a novel perspective for understanding the intra-tumor heterogeneity in HNSCC.

## Discussion

In this study, we present SpaCNA, a spatial-aware framework for CNA detection from spatial transcriptomics data. Given the inherent high noise and sparsity of ST data, as well as the signal dilution caused by the tumor/normal cell mixture, accurate identification of CNAs poses significant challenges. To address these issues, our approach leverages multi-modal spatial information through the construction of spot-neighbor graphs, which enhances the raw expression values and allows for the fitting of a HMRF model and finally assigns CNA states. Through systematic evaluations across a range of simulated scenarios and real-world datasets, SpaCNA consistently achieves high precision and recall. Building on the accurate and reliable detection results, we further introduce downstream modules that estimate the malignancy levels and boundary scores for each spot, thereby expanding the applicability of SpaCNA in the analysis of tumor ST data.

In the design of SpaCNA algorithm, we primarily rely on the similarity of spots. Similar to many previous studies on clustering or spatial domain detection, we naturally hypothesize that adjacent spots exhibit similar states. However, this assumption may not be suitable for spots located at the tumor boundaries. To achieve a more accurate similarity assumption, we extract local image feature from the H&E stained image and exclude adjacent spots that exhibit significant histological differences. In practical, H&E stained images are not always available. Fortunately, the impact at the boundaries is not significant, allowing us to omit this step in certain samples while still achieving reliable results. After constructing the adjacency graph, we utilize the similarity assumption in two aspects. In the step of enhancing gene expression, we explicitly use weighted average to amplify the signal of individual spots, ensuring that weaker CNA signals are not overlooked. In the state assignment step, we implicitly constrain the hidden states of similar spots to be more likely identical through the HMRF model, resulting in a final outcome with reduced spatial noise.

We leveraged the results of SpaCNA to analyze the tumor microenvironment across various tumor types. In a breast cancer slice, we delineated the tumor core and boundary based on the detected CNAs, which closely aligned with the staining patterns. The boundary spots were enriched with pathways and cell types associated with the cell cycle, exhibiting significant differences from those in the core region. In a CRC slice, we identified two distinct subclones through CNA clustering. One subclone exhibited a lower proportion of tumor cells and a higher presence of immune cells at the boundary, while the other clone displayed a contrasting profile. In these analyses, CNA primarily serves to define the structure of tumors. Compared to manual annotations or using expression of individual marker genes, classification based on large CNA segments is more convenient and robust. This is also why there has been a frequent emphasis on inferring CNA in scRNA-seq studies of tumors. In this context, the downstream analysis module we developed enables a more straightforward interpretation of tumor structure in space and can be integrated with various subsequent analyses, making it an effective tool for elucidating the mechanisms of tumor formation and progression.

Additionally, we extended the application of SpaCNA to a novel 3D ST technology. In 19 consecutive slices obtained from Open-ST, SpaCNA identified three spatially coherent subclones that exhibited consistent CNA patterns across all slices. Notably, we identified a clone-specific copy number amplification on chromosome 3q, harboring critical cancer genes such as *PODXL2* and *PIK3CA*, which were closely associated with the prognosis of EGFR-related therapies. The spots carrying this CNA are limited, and other methods, including inferCNV and STARCH, have failed to detect it. Here, the key aspect lies in the construction of the adjacency graph. After aligning the coordinates of all the slices, we establish neighbor relationships among adjacent slices, allowing the CNA of all spots to be incorporated into the model and inferred simultaneously. For other similar multi-slice 3D technologies, as long as the alignment of the slices is available, SpaCNA can be similarly applied and is expected to achieve better results than single-slice analyses.

In recent years, the technology of spatial omics has been continuously iterating and advancing, presenting both new opportunities and challenges for CNA detection. A notable trend is the ongoing improvement in sequencing resolution and throughput^34^. In our analysis of HNSCC, we merged ~1,000,000 single cells into ~20,000 pseudo spots to save computational resources. While this approach sacrifices some detection resolution, it ensures that major CNAs are detectable. Moreover, some new technologies can capture not only the transcriptome but also simultaneously measure other modalities like epigenome^35^. In these types of data, balancing and accommodating multiple modalities poses an additional challenge in designing CNA detection algorithms. The model architecture of SpaCNA has the potential to be applicable to such data. By treating CNA as a common hidden state and the signals of each modality as observed values under different branches, we can establish a similar HMRF model and achieve balance among the modalities by adjusting the weights. Overall, SpaCNA has shown promising performance with existing spatial transcriptomics data, and we hope it will play a broader role in future tumor research.

## Methods

### The SpaCNA method

#### Expression enhancement

##### Introduction

The inputs of the method include the gene-by-spot expression matrix, ***E*** ∈ ℝ^*p×n*^,the spot coordinate matrix ***L*** ∈ ℝ^*n×*2^, and the H&E stained image ***I***. Let *n* denote the number of spots and *p* the number of genes. Due to the high noise and sparsity, we perform smoothing and denoising on ***E*** before the CNA calling to enhance the raw data. Specifically, we first perform spot-wise smoothing based on the order of genes. Then, we construct a spot neighbor graph based on the spot coordinate and H&E stained image. We utilize both expression similarity and histological similarity to further enhance the expression.

##### Spot-wise smoothing

We filter out genes that are expressed in less than 5% of the spots. We then perform library size normalization, converting the raw expression values to counts per million. After log transformation of *E*, we fit a first-order dynamic linear model for each column (i.e., each spot):

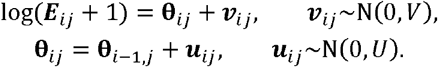

Model estimation is implemented using the R package dlm^36^. Here, we assume that genes are arranged according to their positions on the chromosome. Empirically, we set the parameter *V* to 0.01 and *U* to 0.01. Subsequently, the elements ***E***_*ij*_ of the matrix are replaced with 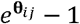, and we still denote the matrix as ***E***.

##### Spot graph construction

At this step, we construct the neighborhood relationships between spots. We aim to obtain an adjacency graph where each point corresponds to a sequencing spot, and two connected spots are likely to exhibit similar expression patterns. We calculate the Euclidean distance between two spots using ***L*** as follows:

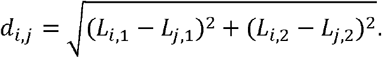

We use the set 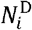 to represent the top 6 spots with the minimum Euclidean distance to spot *i*. We further consider measuring histological similarity using the H&E stained image. For spot *i*, we take a local image tile *I*_*i*_of 224*224 pixels centered at the coordinates of *i*, then extract tile feature *y*_*i*_ *= f*(*I*_*i*_) using pre-trained ResNet-50 model ^37^. Then we calculate Pearson’s correlation between *y*_*i*_ and *y*_*j*_ to measure image similarity:

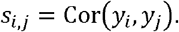

Finally, we only connect two spots when their coordinate is close, and their image similarity is high:

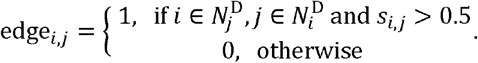

The resulting graph is denoted as **G**, and the neighbors of spot *i* on the graph is denoted as set 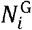. In this way, we aggregated all the spatial information in graph **G**.

##### Smoothing from similar spots

After obtaining the adjacency graph, we enhance by performing a weighted average on similar spots. Here, we also directly consider spots with similar expression profiles. We perform PCA dimensionality reduction on, then calculate pairwise Pearson’s correlation on PCA results. We use set 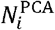 to represent the top 6 spots with the highest correlation to spot *i*. We also assume that spots that are histologically close should exhibit similar expression patterns. Finally, we use both 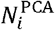 and 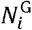 to perform enhancement:

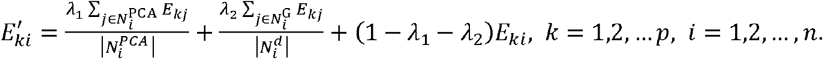

Here, *λ*_1_ and *λ*_2_ are tunable parameters, and both are set to 0.25 by default.

#### Normalization and segmentation

To reduce the computational complexity and perform chromosome segmentation, we divide the chromosomes into non-overlapping bins of fixed lengths (1 Mbps by default). We then merge the gene-by-spot matrix into a bin-by-spot matrix ***X*** ∈ *ℝ*^*m*×*n*^ (*m* is the number of bin). The elements of this matrix can be considered as the read depth within each bin. If a bin has a value of 0 in more than 80% of the spots, or if the number of uniquely mappable positions within the bin is less than 50%, we exclude that bin from ***X***.

To provide an initial estimation of the copy ratio, we need to pre-identify a subset of non-tumor spots. Based on this subset, we can establish a reference read depth baseline in the absence of CNAs. In practice, we first perform spot clustering using the default workflow of Seurat^38^. Then, we manually extract the cluster with the lowest expression values of known tumor marker genes. After extracting the corresponding columns from ***X*** for these spots, we calculate the median for each row to obtain the baseline expression level of the genes, representing the expression level in the absence of CNA. From this baseline vector ***x***^normal^ and ***X***, we obtaina copy number ratio matrix ***R*** as follows:

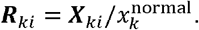

For the subsequent inference of copy number states, we merge adjacent bins into segments. Here, we adopt the multi-sample BIC-seq algorithm for segmentation. Based on the bin read counts of tumor and normal genomes, the algorithm uses the BIC criterion to merge bins iteratively and report the final segmentation result. We use it to determine common breakpoints of all spots from the read depth matrix ***X*** and the baseline ***x***^normal^. Specifically, we denote the segmentation of bins as a vector S = (*s*_0_, *s*_1_, …, *s*_L−1_, *s*_*L*_), where *s*_0_ = 0 < *s*_1_ < … < *s*_*L*− 1_ < *s*_*L*_ = *m* represent the positions of breakpoints and the *l*-th segment consists of consecutive bins from *s*_*i*−L_ + 1 to *s*_*l*_. The algorithm minimizes the following objective function:

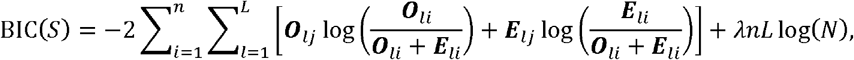

where 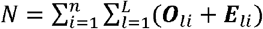,***O***_*li*_ represents the total observed read depths of *i*-th cell in *l*-th segment and ***E***_*li*_ represents the total expected read depths when no CAN occurs. They are calculated from ***X*** and ***x***^normal^:

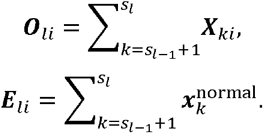

Tuning parameter *λ* is set to 5 by default. We obtain the optimal segmentation 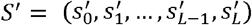 from BIC-seq algorithm.

#### Copy number state inference

##### HMRF model

We use a Hidden Markov Random Field (HMRF) to model the relationship between the copy number states and the copy number ratio, while incorporating spatial similarity. In the model, the copy number ratio ***R*** obtained in the previous step serves as the observed values, while the true copy number states are the hidden states. We assume that bins within the same segment share same copy number state. We denote these hidden states by a segment-by-spot matrix ***S*** ∈ ℝ^*L*×*n*^, where each element is one of {*gain,loss,neutral*}.

We independently model and estimate the hidden states for each segment. Based on the common settings of HMRF, we assume that the joint distribution of segment *l* is as follows:

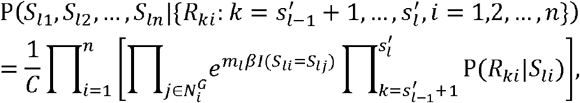

where *C* is the scale factor and 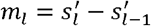 is the length of segment. In the above equation, the first term represents the spatial constraint, with the tunable parameter *β* controlling the strength of this constraint. It intuitively encourages neighboring spots to have the same copy number state. The second term is the emission probability of the observed values given the hidden states. We assume that (· | *s*) follows Gaussian distribution 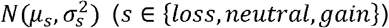. Let ***μ*** = (*μ*_*loss*_, *μ*_*neutral*_, *μ*_*gain*_),***σ*** = (*σ*_*loss*_, *σ*_*neutral*_, *σ*_*gain*_), the likelihood function can be written as:

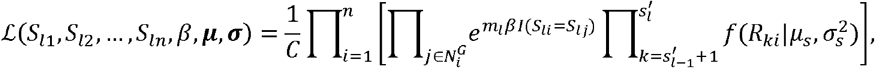

where 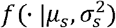 is the density function of 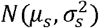.

##### Parameter initialization

We use simulated data to determine *μ*_*s*_ and 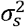. In general, we fit the distribution of each gene using the raw expression matrix ***E***, then simulate an expression matrix with given CNAs by adjusting the means of these distributions. We then apply the same normalization step to obtain the copy number ratios from the simulated data and estimate P (· |*s*) based on the CNAs we set.

In previous studies, it is commonly assumed that gene expression follows a zero-inflated negative binomial distribution (ZINB). The probability density function of ZINB is given as follows:

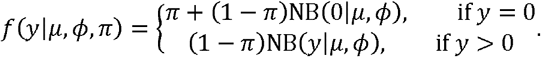

NB (· | *μ, ϕ*) denotes the negative binomial distribution with a mean parameter *μ* and a dispersion parameter *ϕ*, and *π* represents the zero-inflation parameter.

We first define the distribution of each gene *t* (*t* = 1, 2, …, *p*) under normal copy number states as ZINB(*μ*_*t*_, *ϕ*_*t*_, *π*_*t*_). Since there is no analytical form for ZINB parameter estimation, we adopt a simple approach to approximate all the parameters. We calculate the mean of each row in ***E***^0^ and the proportion of zero elements, denoting them as the vectors ***e***^0^ and ***π***^0^, respectively. We directly use ***e***^0^ as the mean parameters. We then assign a fixed value for the dispersion parameter of all genes, denoted as *ϕ*_0_ (defaulting to 0.1). For the zero-inflation rate, considering that the proportion of zero elements for individual genes are quite unstable, we learn the relationship between ***π***^0^ and ***e***^0^and use the fitted values as a substitute. Specifically, we use the spline function *smooth*.*spline()* in R to fit the relationship between ***π***^0^ and log***e***^0^:

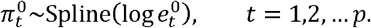

Based on the above results, we set 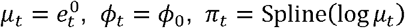, resulting in the expression distribution for gene *t* as ZINB(*μ*_*t*_, *ϕ*_*t*_, *π*_*t*_). Correspondingly, if gene *t* harbors a copy number gain at certain spots, its distribution changes to ZINB (1.5 *μ*_*t*_, *ϕ*_*t*_,Spline (log 1.5 *μ*_*t*_). If there is a copy number loss, the distribution becomes ZINB (0.5 *μ*_*t*_, *ϕ*_*t*_,Spline (log 0.5 *μ*_*t*_). For each gene, we obtain three distributions, denoted as 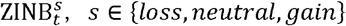.

Next, we simulate the expression matrices. We first simulate the expression profiles of *n*′ = 100 normal spots using 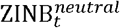. We independently sample *n*′ values from each distribution and synthesized the simulated expression matrix 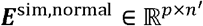. To simulate expressions for tumor spots, we randomly selected 4 chromosomes as regions of copy number gain and another 4 chromosomes as regions of copy number loss. For the genes located in the former, we extracted expression values from the corresponding 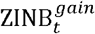, while for the genes in the latter, we generated values from 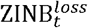. This also yielded a simulated expression matrix 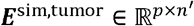.

We combine ***E***^sim,normal^ and ***E***^sim,tumor^ into a single matrix 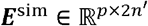, from which we derive the copy number ratio matrix ***R***^*sim*^ following the normalization process previously described. In this approach, we do not simulate spatial coordinates and adjacency graphs, thereby omitting the related steps involved in gene expression enhancement. Furthermore, since the expression levels of normal spots are already established, the baseline expression levels are constructed directly from ***E***^sim,normal^. Finally, the initial values of *μ*_*s*_ and 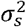 are determined by calculating the mean and variance of expression for regions characterized by copy number gains, losses, and those considered normal.

An initial estimation of *S*_*li*_ can also be obtained by maximizing ℒ without incorporating the spatial constraint term:

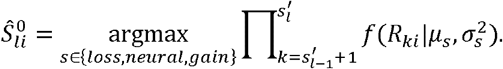

##### Copy state inference

We adopt a local iterative approach to greedily maximize the likelihood function and estimate *S*_*li*_. At each iteration, we update the state of one spot while fixing the states of all other spots:

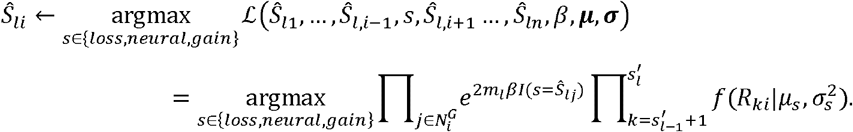

We began by using 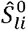 as the initial estimate. In each round of iteration, we update all spots one by one in a random order, following the method described above.

We also update ***μ*** and ***σ*** at the end of each round. The maximum likelihood estimates for *μ* _*s*_ and 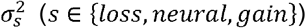 are obtained as follows:

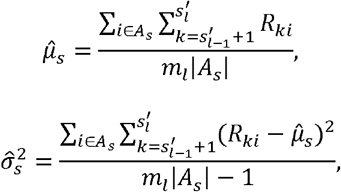

where *A*_*s*_ = {*i* : *Ŝ*_*li*_ = *s*} represents all spots whose current copy number state is estimated to be. After each round of iteration, we update the parameters to the above MLE only if | *A*_*s*_ | >10.

The update is terminated when the maximum number of rounds (5 by default) is reached, or no changes occur after a full round. We found that the iteration usually converges after 5 rounds (Supplementary Figure 9). It is evident that all updates described above result in a non-decreasing likelihood function, which guarantees that the algorithm will always converge. However, it is important to note that these iterations do not ensure convergence to a global optimum. Consequently, it is essential to assign reasonable initial values, which is why we compute them using simulated data rather than assigning them randomly.

#### Estimation of malignancy level

In chromosomal regions where CNAs are present, spots with higher tumor content display stronger copy number ratio signals, leading to greater deviations from a ratio of 1. Previous researches^39,40^ have employed copy number ratios in critical genomic regions to assess malignancy, effectively differentiating between tumor and normal loci. These studies assume that specific CNAs are established on certain chromosomes (for instance, glioblastoma multiforme is known to typically exhibit copy number gains on chromosome 7). Therefore, the malignancy level of spot *i* can be estimated as:

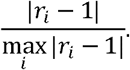

Here, *r*_*i*_ represents the copy number ratio reported by the CNA method. For many tumors, it is not possible to predict in advance which chromosomal regions will contain CNAs. Moreover, estimating malignancy based solely on a single CNA segment can be unstable and may introduce significant bias.

Based on a similar concept, we use the copy number ratio ***R*** and the copy number state ***S*** to estimate the malignancy level of each spot. We first perform hierarchical clustering on ***R*** to classify all spots into classes (7 by default). We assume that each cluster essentially shares the same copy number state. For a given class 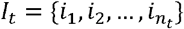 and a segment *k*, if more than 50% of the elements in {*S*_*ki*_:*i*∈*I*_*t*_} are in the same state *s*_*k*_ and *s*_*k*_ *≠ neutral*, we consider that *I*_*t*_ exhibits a consistent CNA in segment *k*. We take into account the set of segments *K*_*gain*_ that exhibit consistent copy number gains and the set *K*_*loss*_ that exhibit consistent copy number losses. Denote the malignancy level of spot *i* as *α*, then the following linear relationship should hold (Supplementary Figure 10):

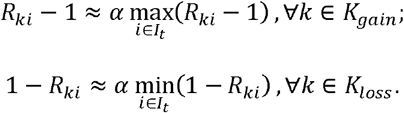

Therefore, we treat *α* as the coefficient in a linear regression model and estimate it using the least squares method. If a particular class exhibits minimal copy number variation, such that | *K*_*gain*_| + |*K*_*loss*_| is below a specified threshold (5 by default), we consider this cluster to be normal spots, assigning a malignancy level of 0. For other clusters, we individually fit the linear regression for each spot to estimate malignancy level. In subsequent analyses, we also interpret this value as indicative of the proportion of tumor cells within the spot.

#### Definition of boundary score

At the boundary between tumor and normal regions, the CNA alteration around spots is greater, which can be effectively captured by our HRMF model. For cell *i* and segment *l*, if we assume that all other latent states are fixed, we can approximately have:

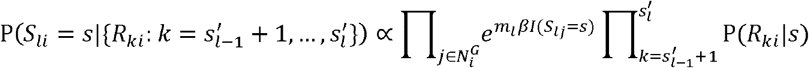

In this discrete posterior distribution, if the probabilities of different copy number states are similar, it indicates that there is a significant variation near the locus, suggesting a higher likelihood of being at the tumor boundary. Normalizing the above expression yields the posterior probability, denoted as 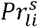. We quantify this uncertainty using entropy, and then sum the entropies of all segments to define the boundary score:

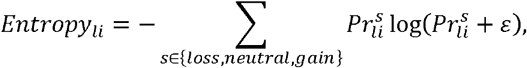

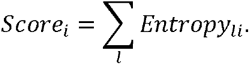

#### Simulation study

##### Generation of simulated data

In the simulation, we assumed that sequencing spots distributed as a 20×20 square grid, resulting in a total of *n =* 400 spots, with each spot directly adjacent to 4 other spots. To simulate the expression matrix, we sampled expression values for each gene from a zero-inflated negative binomial (ZINB) distribution. The mean parameter for the expression distribution at non-tumor spots was obtained from a reference dataset, while for tumor spots, we randomly selected chromosomal regions containing CNAs and modified the mean of the distribution to simulate CNA events. Additionally, we incorporated subclones with different CNA profiles in the simulation, where each subclone contained several spatially clustered spots (specifically, subclone 1 contained the 10×10 region in the bottom-left corner, subclone 2 contained the 8×8 region in the top-right corner, and subclone 3 contained the 8×7 region in the top-left bottom-left corner).

We first simulated the copy number status of each subclone. We divided each of the 22 autosomes into three equal-sized segments and assumed that CNAs occurred at the granularity of these segments. We hypothesized that there were *L*_1_ shared CAN segments among the subclones, and each subclone had an additional *L*_2_ unique CNA segments. We randomly sampled the positions of these CNA segments from the total pool of 66 possible segments and then randomly assigned the alteration type as either copy number gain or loss. We represented the resulting CNA states using a matrix *C* ∈ ℝ^*pxn*^, where *C*_*t,i*_ ∈ {*loss,neutral,gain*} denotes the copy number state at spot *i* for gene *t*.

Next, we set the expression distribution for each gene and then sampled from these distributions to generate the gene expression matrix. We used to ZINB(*μ*_*t*_, *ϕ*_*t*_, *π*_*t*_) to represent the distribution of gene under normal copy number status, where and denote the mean and dispersion parameters of the negative binomial distribution, respectively, and *π*_*t*_ represents the zero-inflation coefficient. We set the mean expression for each gene using a real non-tumor dataset, and then assigned the same *ϕ* and *π* to all genes. By varying the values of *ϕ* and *π*, we could simulate data with different noise levels and missingness rates. Based on the CNA matrix *C*, we independently sampled each element to generate the express ion matrix ***E*** ∈ ℝ^pxn^ as follows:

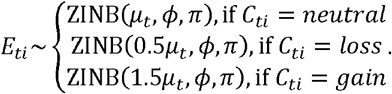

We also considered a more complex scenario, where normal cells were admixed with tumor cells in the tumor sites, leading to a dampening of the CNA signal. To model this, we first sampled the tumor fraction ***α*** ∈ ℝ^*n*^ for all sites from a uniform distribution *U*. For genes without CNA, the generation of *E*_*ti*_ remained the same as before. If, we *C*_*ti*_ = *gain*, we generated the gene expression for the tumor and normal components separately based on *α*_*i*_, and then summed them to obtain *E*_*ti*_:

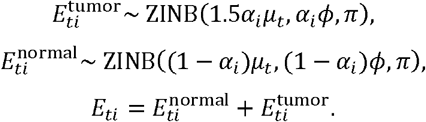

Similarly, if *C*_*ti*_ = *loss*, we used:

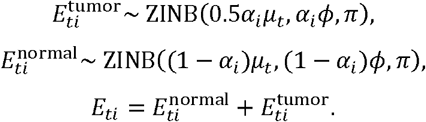

In the above workflow, we varied several parameters to generate data under different simulation settings, including the dispersion parameter *ϕ* (with values {0.1, 0.2, 0.3}), zero-inflation parameter *π* (with values {0.1, 0.2, 0.3}), the number of shared CNA segments among subclones *L*_1_ (with values {3, 5, 7}), and the number of unique CAN segments per subclone *L*_2_ (with values {1, 2}). In the mixture scenario, we also adjusted the distribution *U* of the tumor fraction (with values {*U* (0.65, 0.75), *U* (0.75, 0.85), *U* (0.85, 0.95)}). For each parameter configuration, we generated five replicate datasets and computed various evaluation metrics. In the non-mixture scenario, we primarily investigated the effects of *ϕ* and *π*, and thus averaged the results across different *L*_1_ and *L*_2_ settings. In the mixture scenario, we similarly performed averaging to examine the effects of *ϕ, π* and *U*.

##### Benchmark settings

The simulated expression matrix and normal cell indices are used as input for inferCNV, copyKAT, STARCH, and SpaCNA. For inferCNV, we set the cutoff=0.01, HMM=T, and HMM_type=i3. For copyKAT we set LOW.DR=0.01, while other parameters are kept as default. For STARCH, we include the simulated spatial coordinates as input and set *n_clusters* to the true number of subclones. For SpaCNA, since there is no H&E stained image, we construct the adjacency graph using the four nearest spots within the same clone.

We evaluate each method’s output against the true values (i.e., matrix *C*). SpaCNA, inferCNV, and STARCH all report copy number states (i.e., neutral, gain, and loss). CopyKAT outputs the logarithmic estimates of the copy number ratio, which we convert into these three states using fixed thresholds (values greater than 1.03 are classified as copy number gain, values less than 0.97 as loss, and the remainder as neutral). We then compile the reported results from each method into a matrix and compare it elementwise with *C*. We categorize all elements into the following four types for counting: true positives (copy number gain or loss, correctly classified), false positives (classified as copy number gain or loss but misclassified), true negatives (copy number neutral, correctly classified), and false negatives (copy number gain or loss but classified as copy number neutral). The precision and recall can then be calculated as follows:

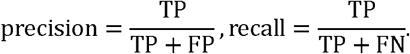

The F1 score is defined as the harmonic mean of precision and recall:

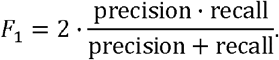

We also compare the accuracy of various methods for clustering. Besides STARCH, we perform hierarchical clustering based on the CNA matrix and categorize all spots into four clusters. STARCH outputs clustering results directly, hence no additional clustering is necessary. Let ***V*** *=* (*v*_*1*_, *v*_*2*_, …, *v*_*n*_) represent the clustering results of a method, and ***U*** =(*u*_*1*_, *u*_*2*_, …, *u*_*n*_) denote the true labels. We utilize the Adjusted Rand Index (ARI) to assess the accuracy of the clustering:

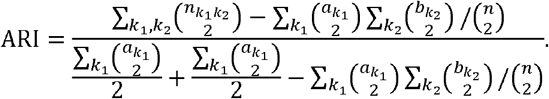

Here,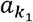 represents the number of spots in ***V*** with label *k*_1_,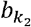 denotes the number of spots in ***U*** with label *k*_2_, and 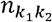 indicates the size of the intersection of loci labeled *k*_1_ in ***V*** and those labeled *k*_2_ in ***U***. The ARI takes values between 0 and 1, with higher values indicating a greater consistency between the clustering results and the true labels.

#### Real data benchmark

##### Comparison based on WES data from liver cancer data

We utilized FACETS to estimate copy numbers from WES data of liver cancer. We provided sequencing data from normal tissue slices of the same patient as the control input, while keeping other parameters at their default settings. Subsequently, we converted the results into copy number states for each bin, which served as the gold standard.

The settings for each CNA method were similar to those in the simulations, although the selection of normal references varied. For the L (leading edge) slices, where a substantial number of normal spots were present in the data, we selected these spots as the reference. In contrast, the T (tumor core) slices contained almost no normal spots. Therefore, we used N (normal) slices from the same patient. For inferCNV and CopyKAT, we directly merged the two expression matrices and specified that the spots from the N samples were normal. For STARCH, we also combined the spot coordinate matrices and ensured they did not overlap. In SpaCNA, we enhanced the expression matrix of the N slices using the same workflow, then computed the mean for each row to determine the baseline expression levels of the genes, substituting this value for ***x***^*normal*^. Consequently, we calculated the mode of the single-cell copy number states for each chromosomal bin to obtain bulk-level results. This allows us to compare the results with the gold standard provided by FACETS. Similar to the simulation, we calculated precision, recall, and F1 score as evaluation metrics.

##### Accuracy of SpaCNA in Distinguishing Tumor from Non-Tumor

We first estimated the tumor proportion at each locus using the deconvolution method SpaCET and combined H&E stained images to manually annotate tumor spots (Supplementary Figure 11). Then we defined the malignancy level of each spot based on the CNA results from different methods. For SpaCNA, we employed the linear regression approach previously described. For the other methods, we calculated the proportion of chromosomal regions exhibiting CNAs at each spot, utilizing this proportion as a score indicative of malignancy level. For all spots within a given sample, we represented the malignancy scores derived from a specific method as the vector ***s*** ∈ ℝ^*n*^ and the manually annotated labels as ***L***∈ ℝ^*n*^. In cases where the CNA detection results align with the annotations, we would generally expect the malignancy scores for tumor spots to be higher, while those for non-tumor spots should be lower. Consequently, we treated this as a binary classification problem and utilized the area under the receiver operating characteristic curve (ROC AUC) to evaluate the effectiveness of in classifying the true labels ***L***. We also examined whether ***s*** is consistent with the tumor proportions inferred by SpaCNA. We calculated the Pearson correlation coefficient between them and the results are presented in Supplementary Figure 5B.

#### Cell type composition analysis on breast cancer

We used an additional breast cancer scRNA-seq dataset as a reference and employed cell2location to estimate the cell type composition within each spot. This resulted in a spot-by-cell type matrix. We then applied the Wilcoxon rank-sum test to identify differentially enriched cell types between two groups of spots. Specifically, we conducted tests comparing the tumor boundary to the tumor core, as well as the tumor boundary to non-tumor regions, and then identified cell types that were simultaneously significant (p-value < 0.05).

#### Cell type composition analysis on colorectal cancer

We directly utilized the deconvolution results provided by the original study^41^. Two types of cell type annotations are provided: a more general classification of major cell types and a relatively detailed classification of minor cell types. Among these, CMS2 in Figure 4I is one of the minor cell types and each row in Figure 4L represents a major cell type.

To analyze the association between CNAs and cell types across multiple samples, we first performed CNA detection on five CRC slices using SpaCNA. We then extracted the tumor spots for each slice and combined the CNA results to a spot-by-bin CNA matrix ***X***. We similarly combined the deconvolution results to form a spot-by-cell type matrix ***Y***. For each row of ***X*** and each row of ***Y***, we grouped the spots based on copy number states using the former, and then applied the Kruskal-Wallis test^42^ to assess whether there are differences in cell type composition between the groups. In this way, we obtained a p-value for each bin and each cell type, which is presented in Figure 4L.

#### Cancer hallmark pathway enrichment analysis

Using gene expressions, we applied GSVA^25^ to calculate gene set enrichment score for 50 cancer hallmark pathways from MSigDB^43^. We obtained 50 scores for each spot independently and applied Wilcoxon’s rank sum test to detect differentially enriched gene sets between two spot groups. P-values were adjusted by Benjamini-Hochberg’s method.

#### 3D dataset analysis

##### Application of SpaCNA

For the OpenST metastatic HNSCC datasets, we constructed pseudo-spots for each tissue slice by partitioning the x-y plane into non-overlapping grids at a fixed interval 140 units, which balances spatial resolution with data sparsity. Each grid point was treated as the center of a pseudo-spot. For each pseudo-spot, we defined a square neighborhood centered at that point with a side length equal to the interval, and aggregated the gene expression profiles of all cells located within this neighborhood to represent the transcriptional profile of the pseudo-spot.

Given that the OpenST data were already spatially aligned across slices, we set the interval between adjacent slices to 100 units and constructed a spatial graph by using the x, y, and z coordinates of the pseudo-spots. Specifically, for each pseudo-spot, we identified its seven nearest neighbors in 3D space and established edges connecting them, thereby forming a spatially coherent 3D graph structure. Based on the graph, we applied SpaCNA to infer CNAs, enabling the detection of spatially consistent CNA patterns across the 3D tissue structure.

##### Cell type proportion calculation and its distance-dependent variation

We obtained the cell type annotation from the original study^26^, then calculated cell type proportions for each pseudo-spot as the ratio between the number of cells of a given type and the total number of cells within the spot. For each tumor clone, we identified non-tumor spots and computed their nearest-neighbor distances to a given clone region using the *get*.*knnx()* function (FNN R package). We then extracted the cell type proportions of these spots and plotted them against the nearest-neighbor distances. We visualized them using scatter points and LOESS-smoothed curves.

##### Other analysis

In the analysis of T cell subtypes, we utilized the *load*.*reference*.*map()* function from the ProjecTILs R package to retrieve a pre-constructed T cell reference atlas. The UMAP embeddings and associated T cell subtypes annotations were provided by the reference. Subsequently, we applied the *Run*.*ProjecTILs()* function to project the query OpenST spatial transcriptomics data onto the reference atlas, enabling the annotation of T cell subtypes.

Cell-cell communication analysis was performed using the COMMOT Python package, and the ligand-receptor pairs were obtained using the *commot*.*pp*.*ligand_receptor_database()* function with parameters *species=‘human’, signaling_type=‘Secreted Signaling’*, and *database=‘CellChat’*.

Hypoxia score was computed using the *AddModuleScore()* function from the Seurat R package. The hypoxia gene set was obtained from the HALLMARK_HYPOXIA signature in the MSigDB^43^ database.

#### TCGA cohort analysis

To evaluate the prognostic relevance of *EGFR* and *PIK3CA* copy number alterations, we obtained gene-level copy number data (ABSOLUTE, n = 517) from the GDC Hub. Samples were stratified into two groups: (1) those harboring amplification of either *EGFR* or *PIK3CA* alone; (2) those with co-amplification of both *EGFR* and *PIK3CA*. Corresponding clinical survival data (n = 603) were also retrieved from the GDC Hub. Then survival analysis was performed using the R package *survival* and the survival curve of each sample group was estimated using the Kaplan-Meier method. Differences between survival curves were assessed using the log-rank test and a statistically significant difference in overall survival was observed between the two groups (P = 0.0084).

