## Supplementary for "CNA detection from spatial transcriptomics using SpaCNA"

**This supplementary file includes:**

Supplementary Figure 1 to 10

Supplementary Table 1 to 3

**Supplementary Figures**

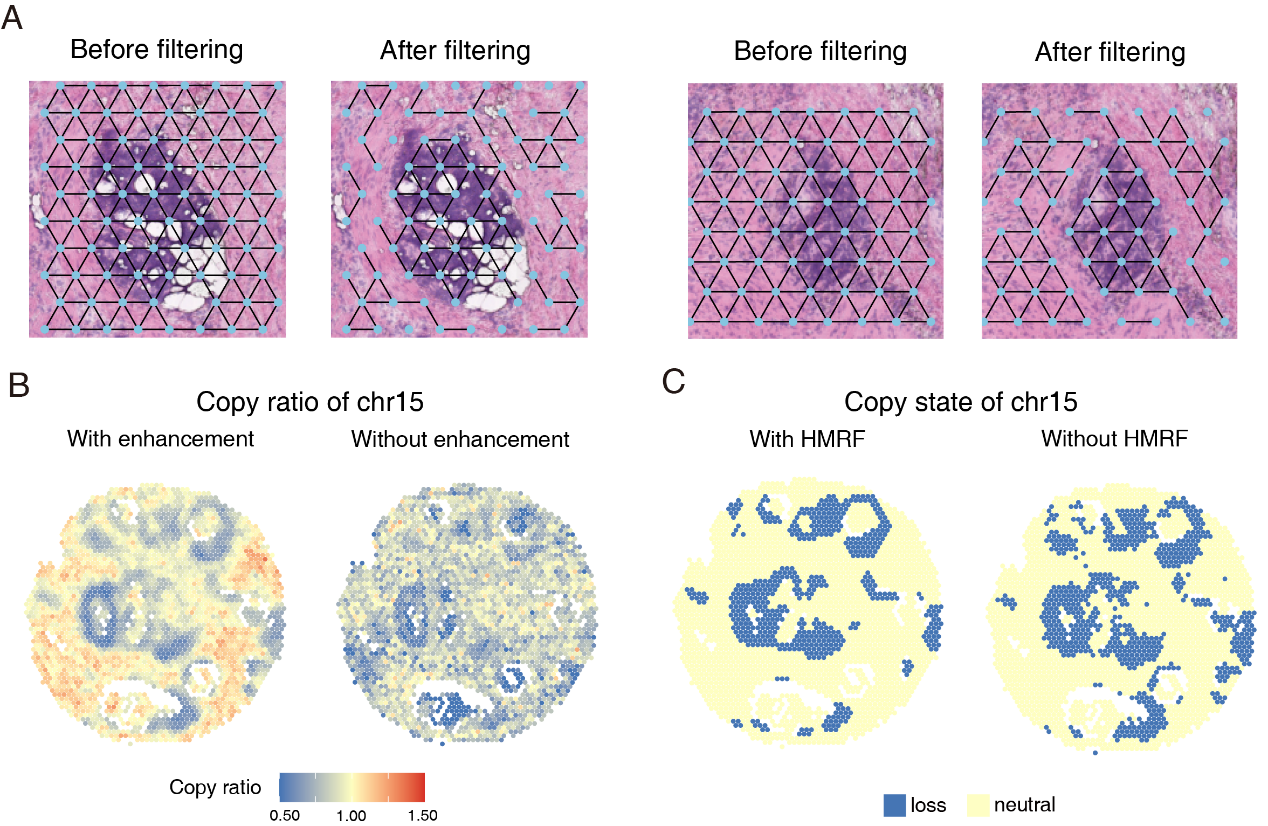

**Supplementary Figure 1. The effects of each step in SpaCNA.** **(A)** The effect of filtering edges using histological features within two local regions. **(B)** The copy number ratio of chromosome 15 calculated by SpaCNA after gene expression enhancement, along with the results obtained by omitting this step in the process. **(C)** The copy number state of chromosome 15 inferred by SpaCNA based on HMRF, as well as the results obtained by removing the spatial constraints from the model.

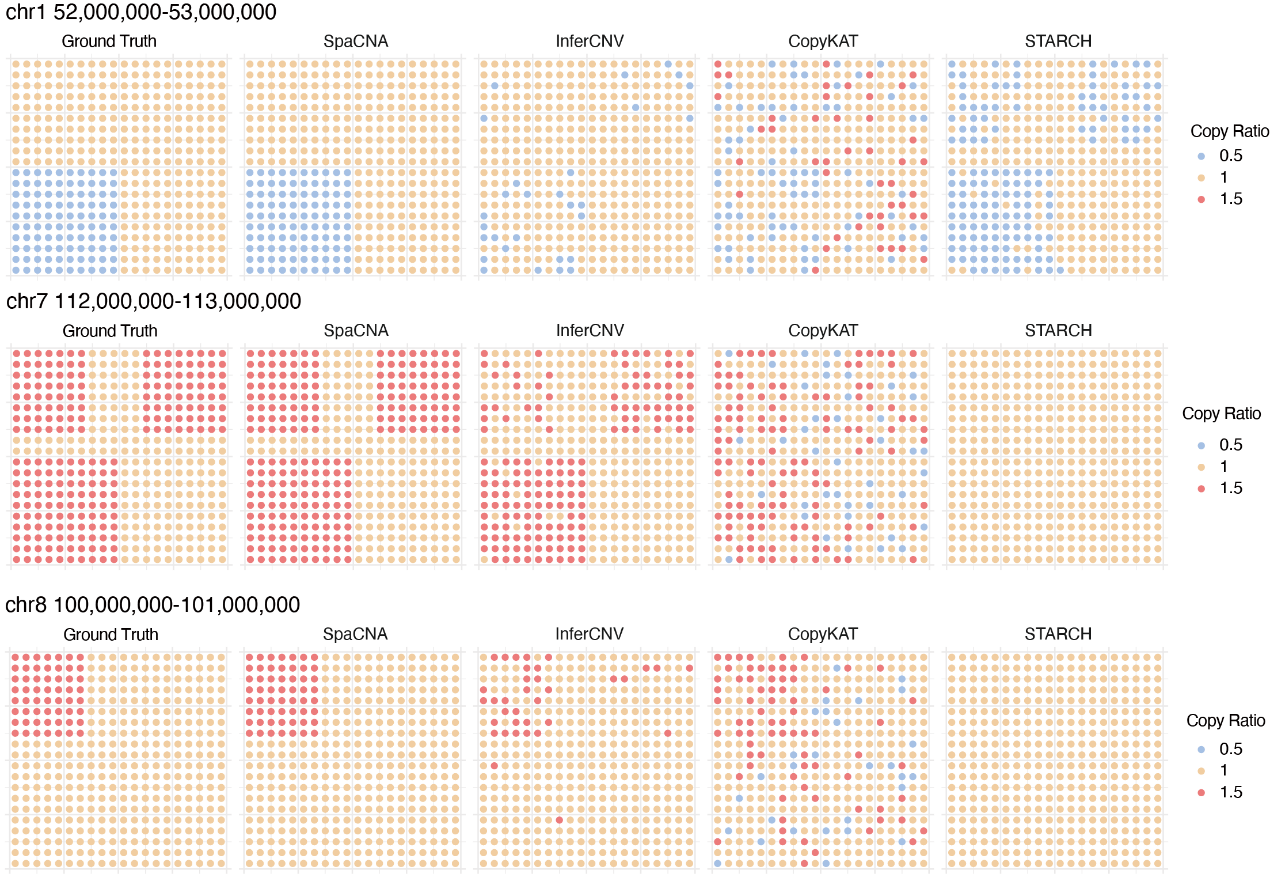

**Supplementary Figure 2. Examples of simulation results.** We used a simulated dataset to demonstrate the effectiveness of CNA detection methods. Each row represents a randomly selected CNA region, and each column represents the ground truth CNA settings and the detection results of various methods.

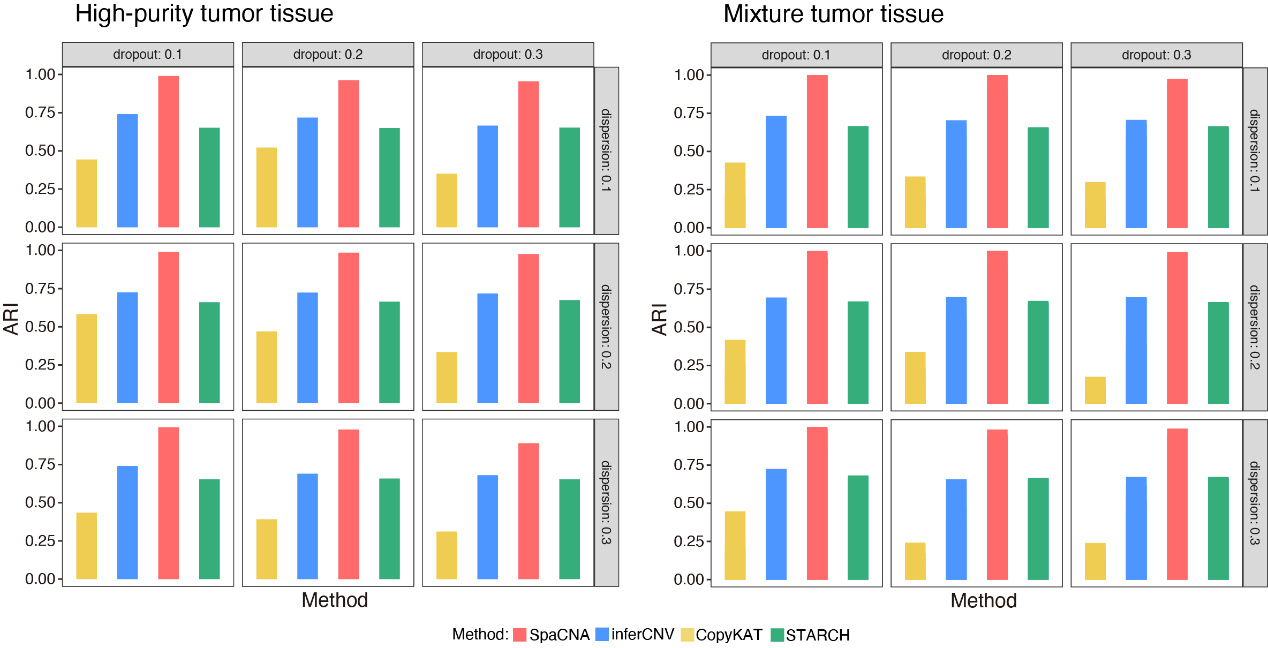

**Supplementary Figure 3. Performance on the simulation study.** The ARI of subclone clustering of SpaCNA and other methods under various dropout and dispersion conditions in the high-purity tumor tissue datasets (left) and mixture tumor tissue datasets (right).

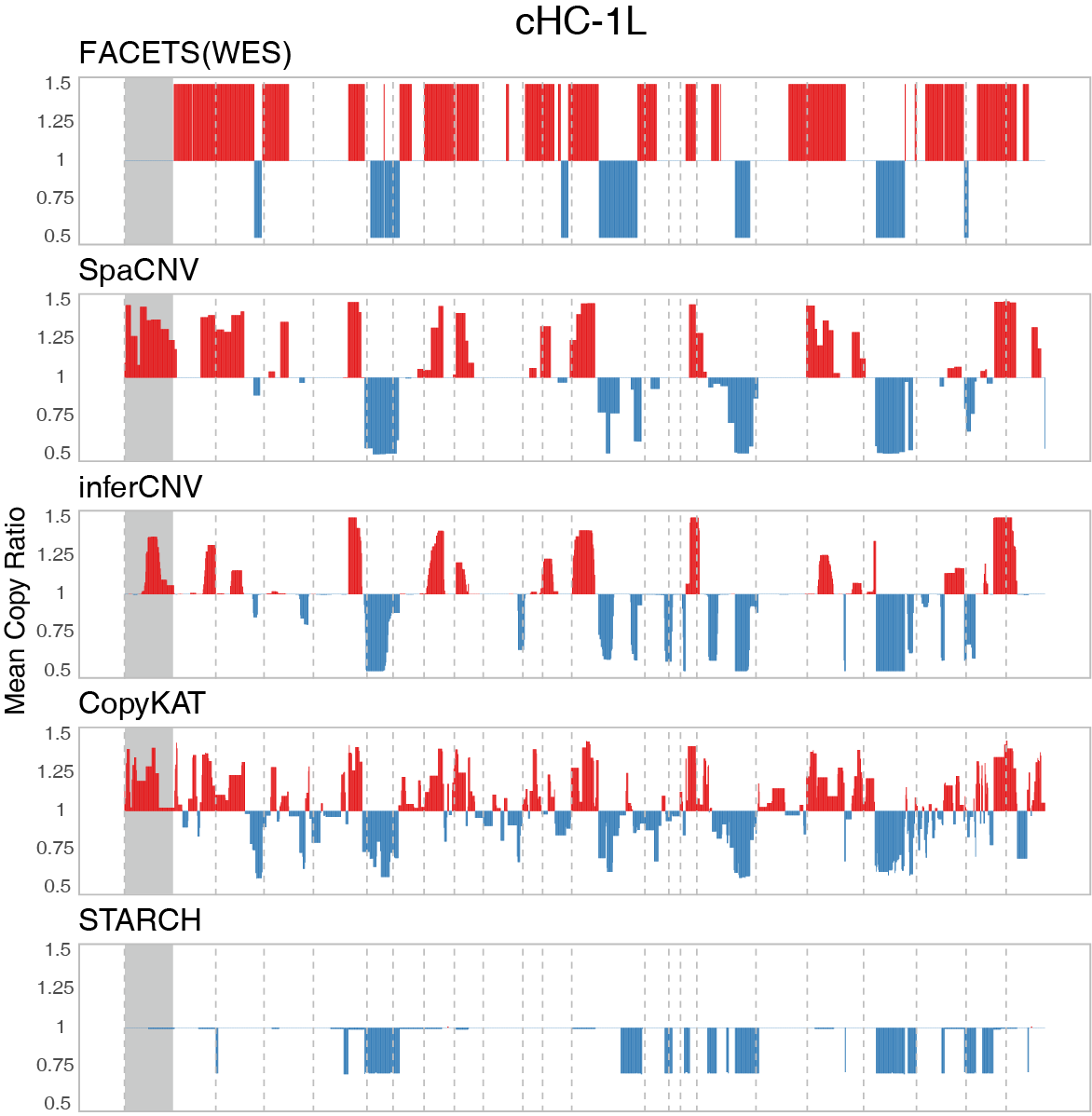

**Supplementary Figure 4. The CNA profiles of cHC-1L.** Each row represents the CNA detection results of a method, with the first row corresponding to the results based on WES. The grey box indicates the chromosome 2p region missed in WES but reported copy number gain by three methods.

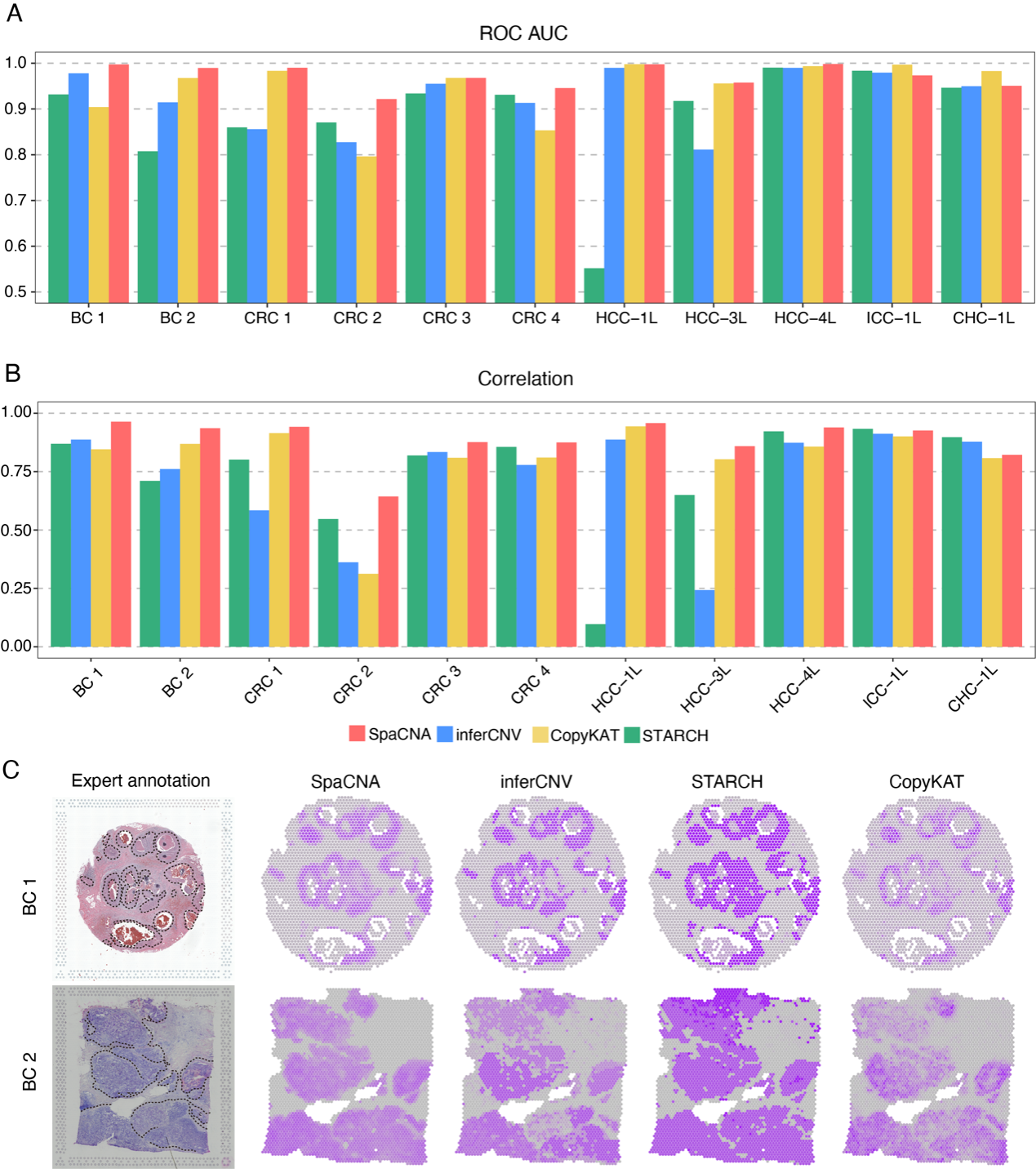

**Supplementary Figure 5. Performance of tumor region identification.** **(A)** The AUROC of four methods in detecting tumor regions. **(B)** The Pearson’s correlation between the malignancy scores and tumor proportions. **(C)** In the first column, the dashed lines on the H&E image indicate the tumor regions annotated by experts, while the remaining four columns represent the malignancy scores calculated from the detected CNA profiles.

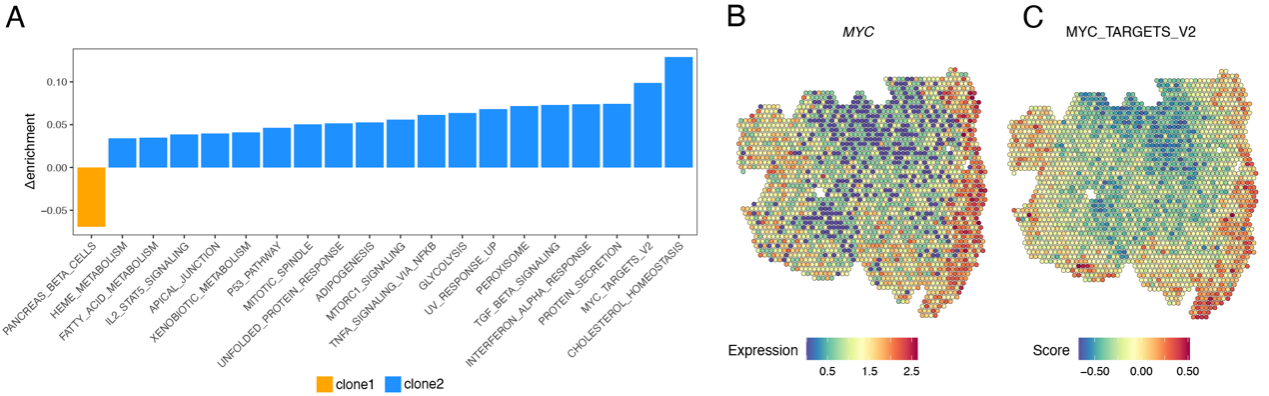

**Supplementary Figure 6. Analysis of CRC sample. (A)** The difference in GSVA enrichment score of cancer hallmark pathways between clone 1 and clone 2. **(B)** The gene expression pattern of *MYC*. **(C)** The MYC_TARGET_V2 pathway enrichment score.

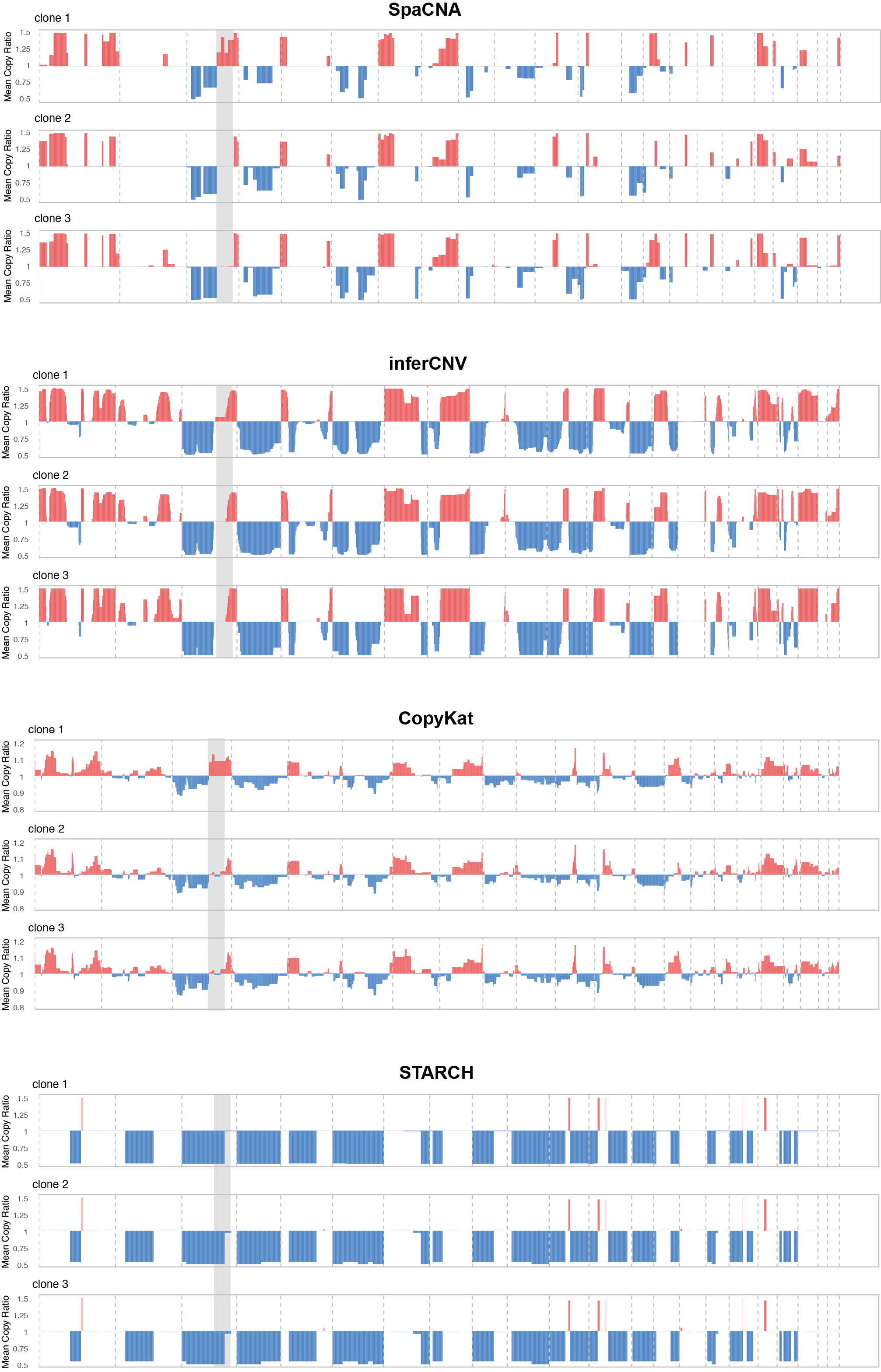

**Supplementary Figure 7.** **The CNA profiles of 3D ST data.** Mean copy ratios for each subclone obtained by SpaCNA and the other methods. The grey boxes indicate a key genomic region located on chromosome 3q. In clone 1, the copy number gain of this region is reported by SpaCNA and CopyKAT, while the other two methods fail to detect it.

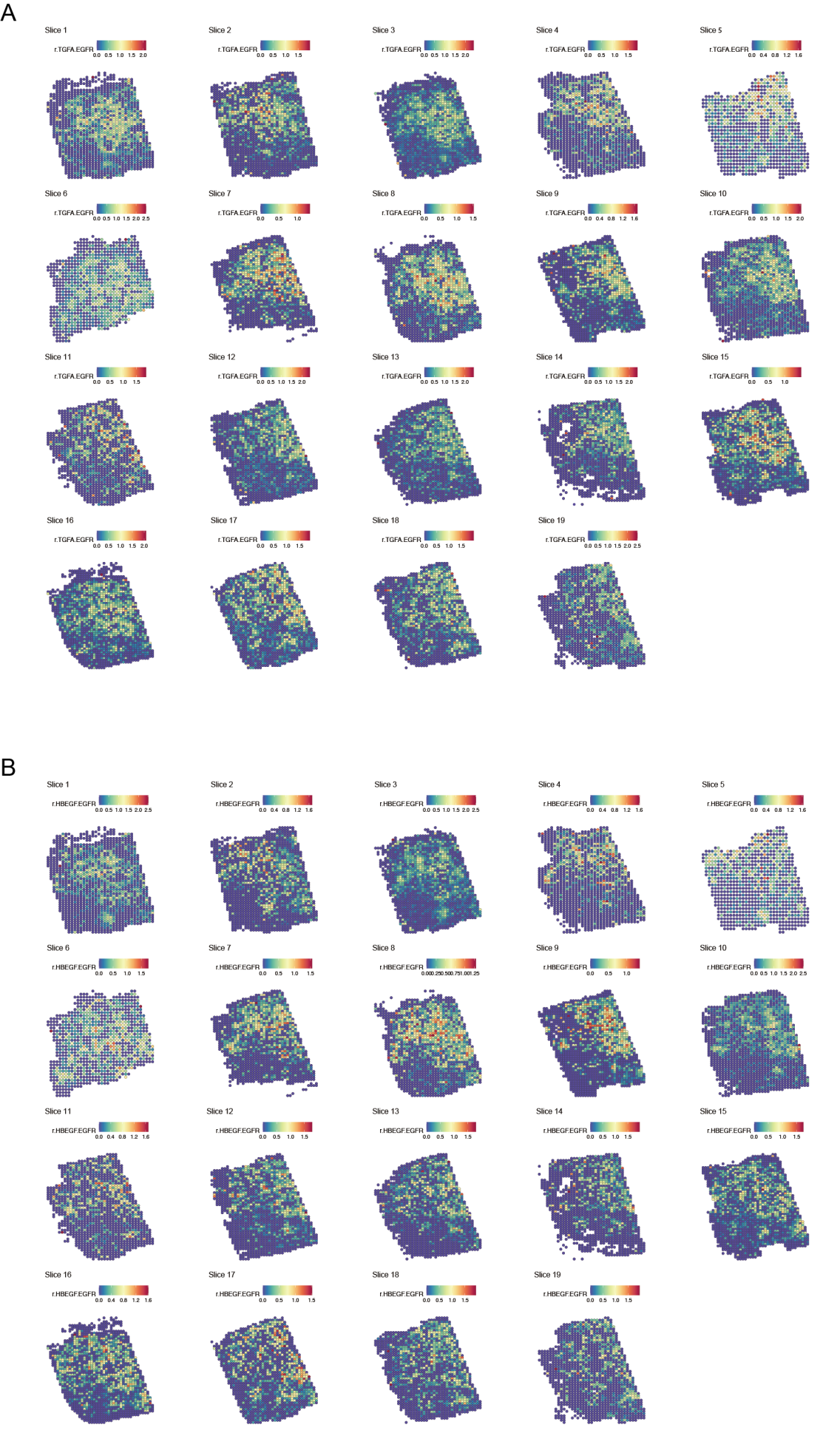

**Supplementary Figure 8. The cell-cell communications analysis of 3D ST data.** The receptor intensity for *TGFA*-*EGFR* (A) and *HBEGF*-*EGFR* (B), as estimated by COMMOT.

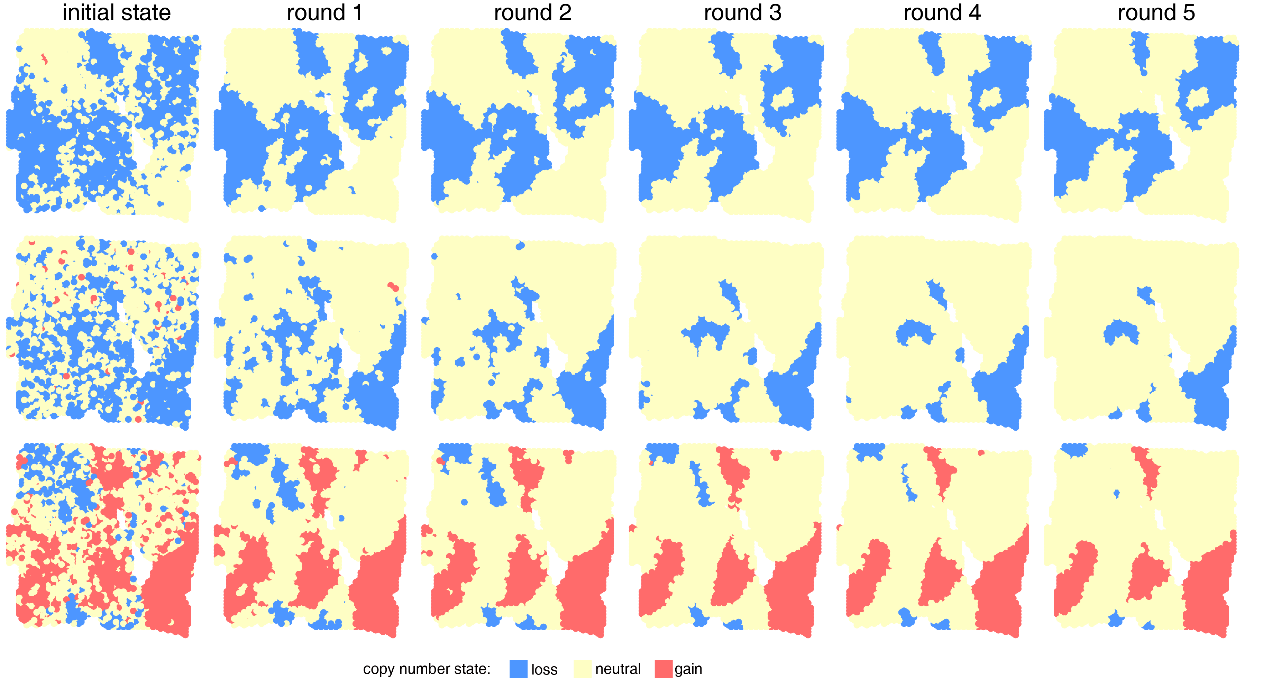

**Supplementary Figure 9. Copy state inference in SpaCNA.** Based on a breast cancer sample, we randomly selected three chromosomal segments and displayed the results of copy number states after each round of iteration. The red, blue, and yellow indicate copy number gain, loss, and neutral, respectively.

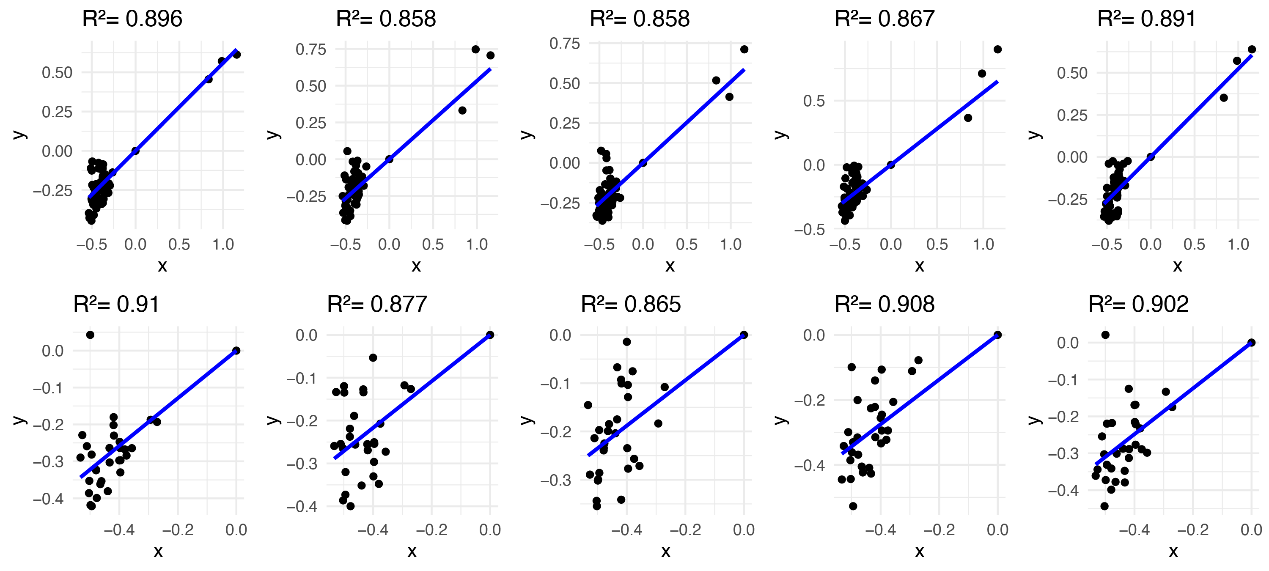

**Supplementary Figure 10.** **Malignancy level estimation in SpaCNA.** Based on a breast cancer sample, we randomly selected 10 spots and then displayed the linear relationship between $R_{ki}-1$ and $\max_{i\in I_{t}} \left( R_{ki}-1 \right)$ ($R_{ki}$ represents the copy ratio of spot *i* in segment *k*). Each point represents a segment, and the x-axis and y-axis correspond to $\max_{i\in I_{t}} \left( R_{ki}-1 \right)$ and $R_{ki}-1$, respectively.

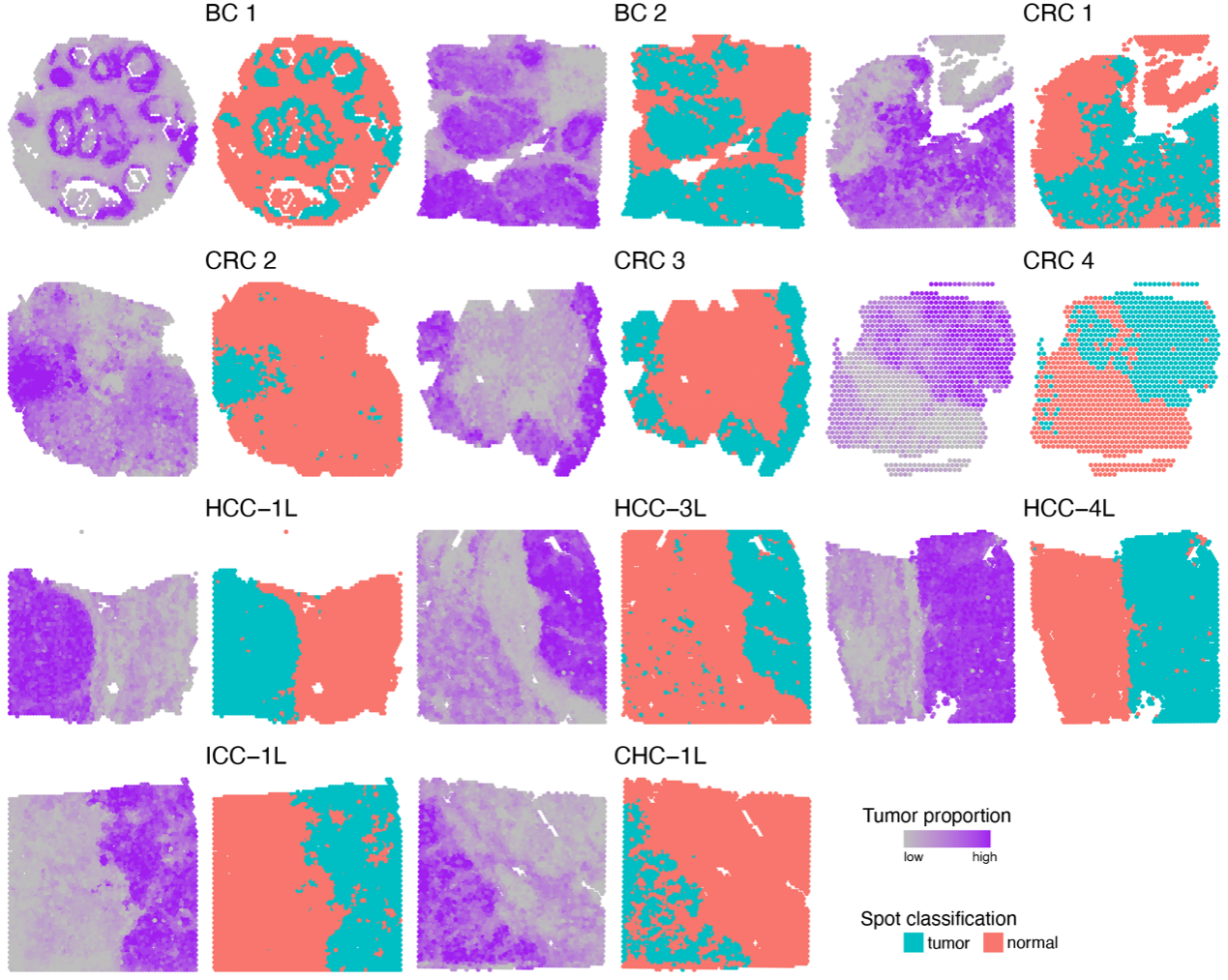

**Supplementary Figure 11.** **Tumor region annotation using SpaCET.** For each ST dataset, we used the deconvolution method SpaCET to estimate the proportion of tumor cells, and then classified the tumor/non-tumor spots using a fixed threshold. The tumor cell proportions (left) and spot classification (right) of 11 datasets are shown.

**Supplementary Tables**

**Supplementary Table 1. Information on liver cancer sections.** Here, N, L, and T represent the normal region, tumor leading edge, and tumor core, respectively. CNA analysis was performed only on the sections from L and T. ICC-1T was excluded due to the presence of a large number of dead cells.

| Section | Type | Spots | Section | Type | Spots |
| --- | --- | --- | --- | --- | --- |
| HCC-1N | N | 2956 | HCC-3N | N | 4289 |
| HCC-1L | L | 2791 | HCC-3L | L | 4758 |
| HCC-1T | T | 3184 | HCC-3T | T | 4456 |
| HCC-2N | N | 4628 | HCC-4N | N | 4397 |
| HCC-2L | L | 4672 | HCC-4L | L | 4113 |
| HCC-2T | T | 4733 | HCC-4T | T | 4162 |
| cHC-1N | N | 2207 | ICC-1L | L | 4654 |
| cHC-1L | L | 4516 |  |  |  |
| cHC-1T | T | 4779 |  |  |  |

**Supplementary Table 2. Precision and recall of CNA detection in L (leading edge) sections.** Bold indicates the highest value in each column.

|  | HCC-1L | HCC-3L | HCC-4L | ICC-1L | cHC-1L |
| --- | --- | --- | --- | --- | --- |
| SpaCNA | **0.74/0.43** | **0.78**/**0.53** | 0.81/**0.59** | **0.60**/0.57 | **0.63**/0.42 |
| inferCNV | 0.73/0.33 | 0.73/0.38 | **0.95**/0.38 | 0.55/**0.58** | 0.45/**0.62** |
| STARCH | 0.51/0.10 | 0.07/0.08 | 0.55/0.16 | 0.56/0.17 | 0.48/0.24 |
| CopyKAT | 0.57/0.43 | 0.16/0.46 | 0.51/0.47 | 0.35/0.45 | 0.37/0.30 |

**Supplementary Table 3. Precision and recall of CNA detection in T (tumor) sections.** Bold indicates the highest value in each column.

|  | HCC-1T | HCC-3T | HCC-4T | cHC-1T |
| --- | --- | --- | --- | --- |
| SpaCNA | 0.78/**0.59** | **0.97**/**0.64** | 0.84/**0.87** | **0.40**/0.66 |
| inferCNV | 0.74/0.55 | 0.97/0.55 | **0.84**/0.83 | 0.35/**0.72** |
| STARCH | 0.66/0.52 | 0.94/0.30 | 0.35/0.50 | 0.53/0.38 |
| CopyKAT | **0.84**/0.27 | 0.93/0.39 | 0.52/0.67 | 0.30/0.52 |
